# Small molecules targeting ARF1 interaction with C9orf72:SMCR8:WDR41 complexes suppress its overactivation implicated in ALS/FTD

**DOI:** 10.64898/2026.01.24.701325

**Authors:** Emma Dixon, Fereshteh Azimian, Angelina Joby Chacko, Rodney Tatum, Christi Boykin, Yan-Hua Chen, Qun Lu

**Affiliations:** Department of Chemistry and Biochemistry, College of Arts and Sciences, University of South Carolina, Columbia, SC, USA; Laboratory of Molecular Neurotherapeutics, Center for Neurotherapeutics, University of South Carolina, Columbia, SC, USA

**Author notes:** Corresponding author: Qun Lu, PhD, MS, BS., Professor of Chemistry and Biochemistry, SmartState Endowed Chair, Center for Neurotherapeutics, Rm 328-329 GSRC, 631 Sumter Street, McCausland College of Arts and Sciences, The University of South Carolina Columbia, SC 29208.

**Keywords:** C9orf72:SMCR8:WDR41, ASAP1, ARF1 small GTPase, Small molecules, Virtual screening, ALS/FTD

## Abstract

The hexanucleotide repeat expansion in *C9orf72* gene is the most common genetic cause of amyotrophic lateral sclerosis (ALS)/frontotemporal dementia (FTD). The C9orf72 protein forms a complex with SMCR8 and WDR41 (CSW), which functions as a GTPase-activating protein (GAP) regulating ARF1 and RAB small GTPases. While these findings implicated ARF1-GAP dysregulation in ALS/FTD and supported ARF1 suppression as potential intervention, small molecules that modulate ARF1-CSW interactions are lacking.

In this study, we demonstrated upregulation of tyrosine-phosphorylated (Tyr-782) ASAP1 (also known as AMAP1, DDEF1, or Centaurin β4), an ARF-GAP, in human motor cortex of both sporadic ALS and ALS with *C9orf72* mutations. Ectopic *C9orf72* expression partially mimicked the effects of a known ARF1 inhibitor brefeldin A to disperse Golgi apparatus. Computer-aided rational drug design with high-throughput *in-silico* screening identified MCULE-5095997944 (Named as SCC944) as a ARF1-CSW modulator. SCC944 binds directly to ARF1 and reduced GTP-bound ARF1 levels upon ARF1 activation. SCC944 demonstrated brefeldin A-like ARF1-dependent alteration of organelle organization including Golgi, microtubules, and mitochondria, but also a protein trafficking pattern that is distinct from brefeldin A mechanism. These studies identified the first small molecule targeting ARF1-CSW interaction and further support ARF1 modulation as a potential therapeutic approach for ALS/FTD.

## Introduction

Amyotrophic lateral sclerosis (ALS) is a fatal neurodegenerative disease that progressively affects motor neurons in the brain and spinal cord, leading to muscle weakness, paralysis, and ultimately death. With an incidence rate of two per 100,000 people, ALS has a similar prevalence to multiple sclerosis, and some patients develop frontotemporal dementia (FTD) (1). Despite decades of intense research, ALS/FTD currently has no cure. Existing treatments, including Riluzole and Edaravone, offer only modest benefits for survival and symptom management (2). Therefore, a better understanding of ALS/FTD pathophysiology is essential for the development of more effective therapeutic strategies.

In familial ALS/FTD, hexanucleotide repeat expansion (HRE) (GGGGCC) in the non-coding region of the *C9orf72* gene has been identified as the most common genetic causes of ALS/FTD (3–5). To combat ALS/FTD with *C9orf72* mutations, neurotoxicity as results of two main pathogenic events must be addressed: (1) the gain-of-function from HRE and the non-canonical expression of the dipeptides versus (2) the loss-of-function *C9orf72* haploinsufficiency. Although considerable effort has been devoted to gene therapy and antisense oligonucleotide (ASO) approaches to counteract HRE-induced toxicity, significant progress is still required in restoring C9orf72 function. The loss of this function can result in serious outcomes, such as impaired cellular waste clearance and increased neuroinflammation.

Small GTPases have been implicated as promising drug targets in neurodegenerative diseases including Alzheimer’s disease (AD), Parkinson’s disease (PD), and ALS (6). The shared common features observed in the pathologies of these diseases include the presence of misfolded protein aggregates and organelle dysfunctions, which are associated with the dysregulation of small GTPases like ARF and RAB (7, 8). Small GTPases transition from an inactive GDP-bound state to an active GTP-bound state through interactions with specific regulators. Guanine nucleotide exchange factors (GEFs) facilitate the release of GDP, allowing GTP to bind in its place, thereby activating the small GTPases. Conversely, GTPase-activating proteins (GAPs) promote the hydrolysis of GTP to GDP, resulting in the inactivation of small GTPases. Additionally, GDP-dissociation inhibitors (GDIs) prevent the dissociation of GDP-bound proteins, thereby maintaining small GTPases in their inactive state (6).

C9orf72 forms a complex with Smith-Magenis chromosome regions 8 (SMCR8) and WD repeat-containing protein 41 (WDR41), which interacts with small GTPases, including ARF1 and several RAB proteins (9–11). The CSW complex acts as a GAP for ARF1, helping to control important cellular activities such as cell proliferation, membrane trafficking, remodeling of the actin cytoskeleton, and cell survival, as well as endolysosome-autophage and mitochondria homeostasis in motor neurons (12, 13). Recent studies suggest that restoring C9orf72 levels or enhancing its function with chemical modulators of vesicle trafficking can promote neuronal survival and mitigate neurodegeneration in mouse models (14, 15). Therefore, small molecules that can directly modulate ARF1-CSW interaction would be a promising strategy for restoring C9orf72 mediated ARF1 signaling in ALS/FTD.

To investigate whether CSW related GAP deficiency can be an underlying cause of ARF1 overactivation in the pathogenesis of ALS/FTD, we first determined that ARF-GAP protein ASAP1 (also known as AMAP1, DDEF1, or Centaurin β4) is inactivated in both sporadic human ALS and ALS with C9orf72 mutations. We then applied rationale design and molecular docking-based virtual screening to identify small molecule modulators (SMMs) targeting ARF1-CSW interactions. Biological validation studies revealed that MCULE-5095997944 (hereafter SCC944) exhibits strong binding affinity for ARF1 with a favorable pharmacokinetics and toxicity profile. Treatment with SCC944 significantly decreased GTP-bound ARF1 levels following its activation and led to reorganization of the ARF1-dependent, brefeldin A-sensitive Golgi, microtubule, and mitochondrial organization. Additionally, our studies identified a protein trafficking pattern altered by SCC944 that is distinct from brefeldin A mechanism. Collectively, these studies establish a pharmacological approach for direct modulation of ARF1 small GTPase activity and support targeting ARF1-CSW interaction as a potential therapeutic strategy for ALS/FTD.

## Results

### ARF1 dysregulation in ALS/FTD

The notion that C9orf72-SMCR8-WDR40 complex acts as an ARF-GAP raised the question whether ARF1 is overactivated in ALS/FTD. Therefore, we first investigated tyrosine phosphorylation state of ASAP1, an ARF-GAP, in posterior frontal motor cortex from human sporadic ALS (sALS) and ALS with *C9orf72* mutations (C9ALS). When ASAP1 is phosphorylated by Src and Pyk2 targeting specific tyrosine residues like Tyr-782, ASAP1 activation is suppressed thereby leaving ARF1 in a sustained GTP-bound form (16). In **Figure 1A**, basal ASAP1 Tyr 782 phosphorylation can be observed in posterior frontal motor cortex of a non-ALS brain. However, the sALS and C9ALS brains demonstrated visibly higher number of positive cells when stained with anti-Phospho-ASAP1 than that in the non-ALS control brains (**Figure 1B and C**). This observation was confirmed through the quantification analysis which showed a significant difference in the percentage of positive cells in control, sALS, and C9ALS brains (**Figure 1D**). About 90% of the cells in sALS and C9ALS motor cortex were Phospho-ASAP1 positive whereas the control brain only showed about 60% positive cells.

**Figure 1.**
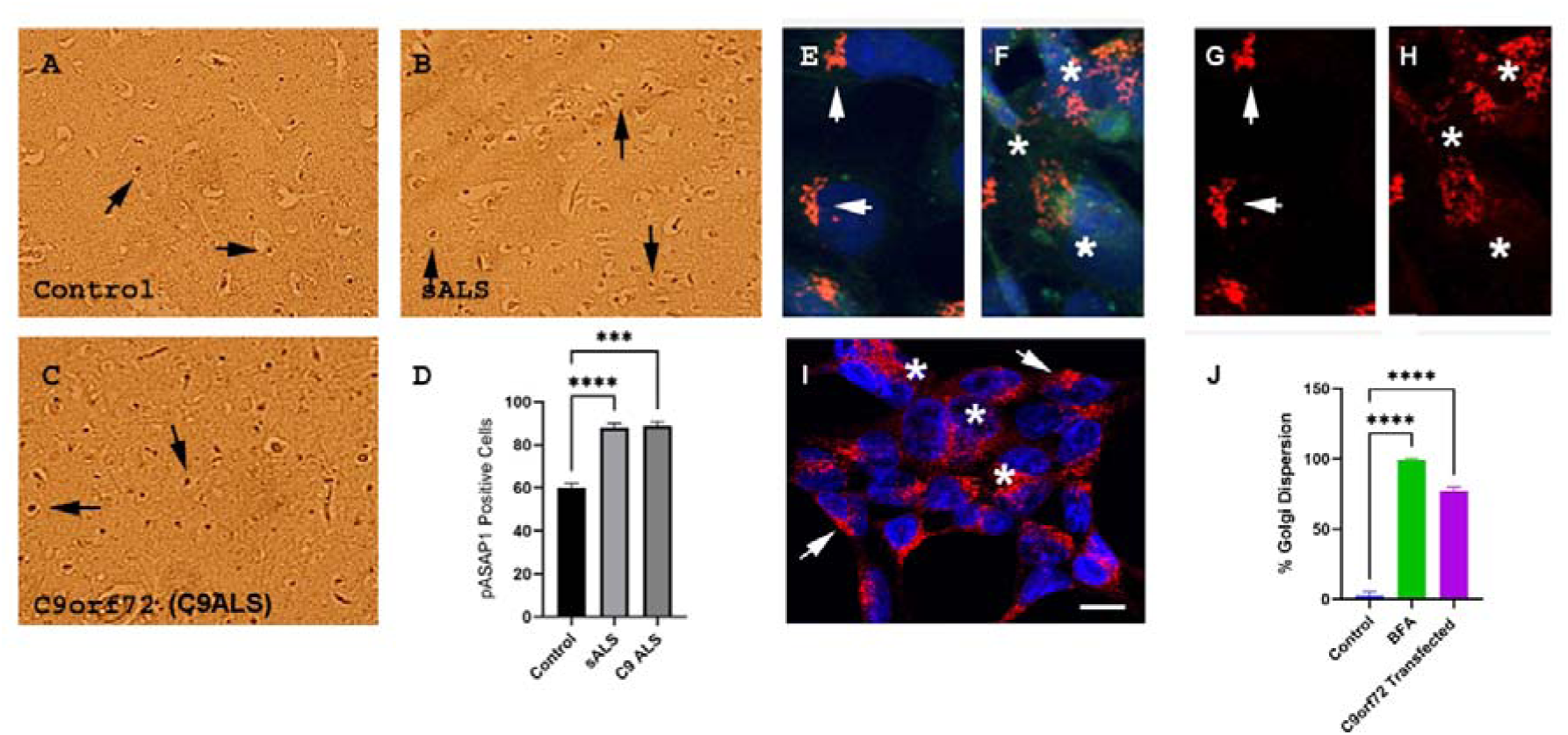
ARF-GAP dysregulation in ALS/FTD. **(A-D)** Elevated Phospho-ASAP1 expression in ALS brains. Expression and comparison of Phospho-ASAP1 (Tyr782) in the posterior frontal motor cortex of **(A)** non-ALS control, **(B)** sALS, and **(C) C9**ALS brain tissue samples. IHC staining for tyrosine phosphorylated ASAP1 was performed in all brain tissue samples. **(D)** Quantification of percent positive cells in non-ALS control, sALS, and C9ALS brain tissue samples. 300x. The mean percent positive cells were compared. \*\*\**P*-value < 0.001; \*\*\*\**P*-value < 0.0001. **(E-J)** Ectopic C9orf72 expression and ARF**1** dependent Golgi organization. HEK293 cells were transfected with *GFP-C9orf72*. Cells were fixed and stained with GM130 antibody (Red). Cell nucleus was stained with DAPI (Blue). **(E)** Merged immunofluorescent images of non-transfected HEK293 cells with GM130 staining. **(F)** Merged immunofluorescent images of *GFP-C9orf72* transfected HEK293 cells with GM130 staining. **(G)** GM130 immunofluorescent staining of Golgi in non-transfected HEK293 cells corresponding to (**E**)**. (H)** GM130 immunofluorescent staining of Golgi in *GFP-C9orf72* transfected HEK293 cells corresponding to (**F**). DAPI was used to stain nuclei (Blue), GFP-C9orf72 (Green), and GM130 positive Golgi (Red). Arrows point to perinuclear localized Golgi. Asterisks point to dispersion of Golgi in the *GFP-C9orf72* transfected HEK293 cells. **(I)** GM130 immunofluorescent staining of Golgi in HEK293 cells treated with BFA. Arrows point to perinuclear localized Golgi. Asterisks point to dispersion of Golgi. Bar: 30 µm. (**J**) Bar graph shows percent dispersion of Golgi apparatus in *GFP-C9orf72* transfected cells in comparison to non-transfected cells or cells treated with BFA. The mean percent Golgi apparatus dispersion was compared. \*\*\*\**P*-value < 0.0001.

As ARF small GTPases regulate intracellular trafficking involving the Golgi organization, we examined the GM130 positive Golgi in HEK293 cells transfected with *GFP-C9orf72* plasmid DNA. HEK293 cells often showed the perinuclear localization of a compact and well-organized Golgi apparatus (**Figure 1E and G, arrows**). Ectopic *C9orf72* expression caused the dispersion of GM130 positive Golgi in the cytoplasm (**Figure 1F and H, asterisks**). Treatment with brefeldin A (BFA), a known ARF inhibitor, led to a more widespread dispersion of Golgi (**Figure 1I, arrows and asterisks; Also see Figure 1J for comparison**). These studies indicated that the pathogenic mechanism of ALS/FTD likely involves the deficiency of ARF-GAP pathway, and the ARF activity is tightly controlled physiologically to homeostatic balances to avoid undue stress on Golgi organization.

### Targeting the interface of ARF1 and CSW complex

To target ARF-GAP pathway that involves C9orf72, we noted that C9orf72 forms a complex with SMCR8 and WDR41, which has been shown to exhibit GAP activity for ARF1 (11, 17). The cryo-EM structure of ARF1-GDP-BeF3 bound to C9orf72:SMCR8:WDR41 (CSW) complex demonstrated that the longin domains of SMCR8 and C9orf72 together create a binding pocket for ARF1, resembling a spoon-bowl configuration (17). Importantly, the Arg^147^ residue in SMCR8 functions as the catalytic Arg finger, being consistent with the characteristic of most GAPs. Additionally, mutations in critical interfacial residues, such as Ile^49^, Gly^50^ of ARF1 and Arg^147^ of SMCR8, significantly reduced or eliminated GAP activity (17). A putative modulator-binding pocket was shown in **Figure 2A**, which surrounds Arg^147^ in the ARF1-CSW complex by including residues of SMCR8, ARF1, and C9orf72 that are within 6.5 Å of Arg^147^. This pocket comprises Ile^46^, Ile^49^, Gly^50^ from ARF1; Asn^64^ and His^65^ from C9orf72; and Ser^110^, Arg^147^, Pro^107^, and Pro^108^ from SMCR8 (**Figure 2B**). Furthermore, we investigated potential interactions between the CSW complex proteins and small GTPase proteins using STRING (<u>S</u>earch <u>T</u>ool for the <u>R</u>etrieval of Interacting <u>G</u>enes/Proteins) to predict protein-protein interactions (**Figure 2C**). The STRING analysis revealed that the CSW complex proteins interact with various proteins implicated in neurodegenerative diseases, including the small GTPases ARF1, RAB11, and RAB39 (**Figure 2C and D**).

**Figure 2.**
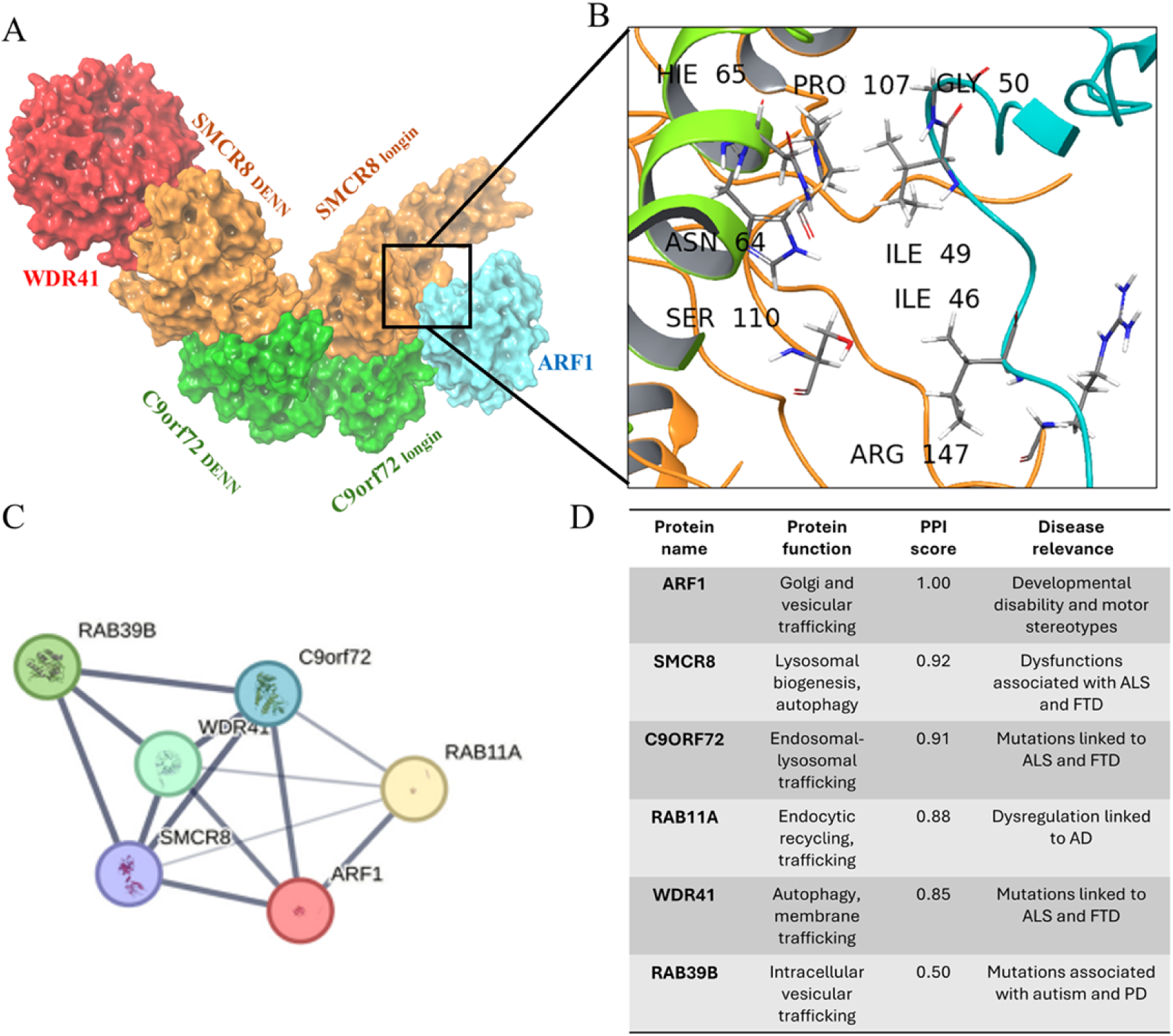
Targeting the interface between ARF1 and CSW complex proteins. **(A)** 3D structural illustration of ARF1-CSW complex and its binding pocket (ARF1 protein is in transparent cyan, SMCR8 and C9orf72 proteins are in orange and green**, respectively** as well as WDR41 is shown in red color (PDB: 7MGE)). The C9orf72 and SMCR8 include both longin and DEN domains. The bracket shows the binding pocket at the interface of ARF1-CSW complex. **(B)** A zoomed-in view of the protein structures highlighting the key residues including the critical Arg^147^ involved in the binding pocket at interface of ARF1 and CSW complex. **(C)** STRING analysis revealed the interaction network of ARF1 and proteins in CSW complex including C9orf72, SMCR8 and WRD41 proteins. Line thickness indicates the strength of data support. **(D)** Table summarizing the functional roles of the proteins in the network, their corresponding interaction scores with ARF1 from STRING analysis, and their involvement in neurodegenerative diseases.

### Structure based hierarchical virtual screening and ADME/Tox profiling

To identify SMMs targeting ARF1-CSW complex, 40,183,714 small molecules in the MCULE library were screened *in silico* according to a rigorous workflow (**Figure 3A**). The top hit compounds demonstrated a variety of interactions with residues at the binding site of ARF1-CSW protein complex. Using Glide module of Schrödinger, the top hit compounds were subjected to sequential HTVS, SP, and XP docking to retrieve 10%, 20%, and 20 % of the compounds in each category, respectively. This progressive screening narrowed down the top candidates to 14 compounds with docking scores ranging from -3.69 to -6.45 kcal/mol (**See Supplementary Table S1**). These SMM compounds follow the favorable Lipinski rule of five with molecular weight below 500 Dalton (**Figure 3A and B**).

**Figure 3.**
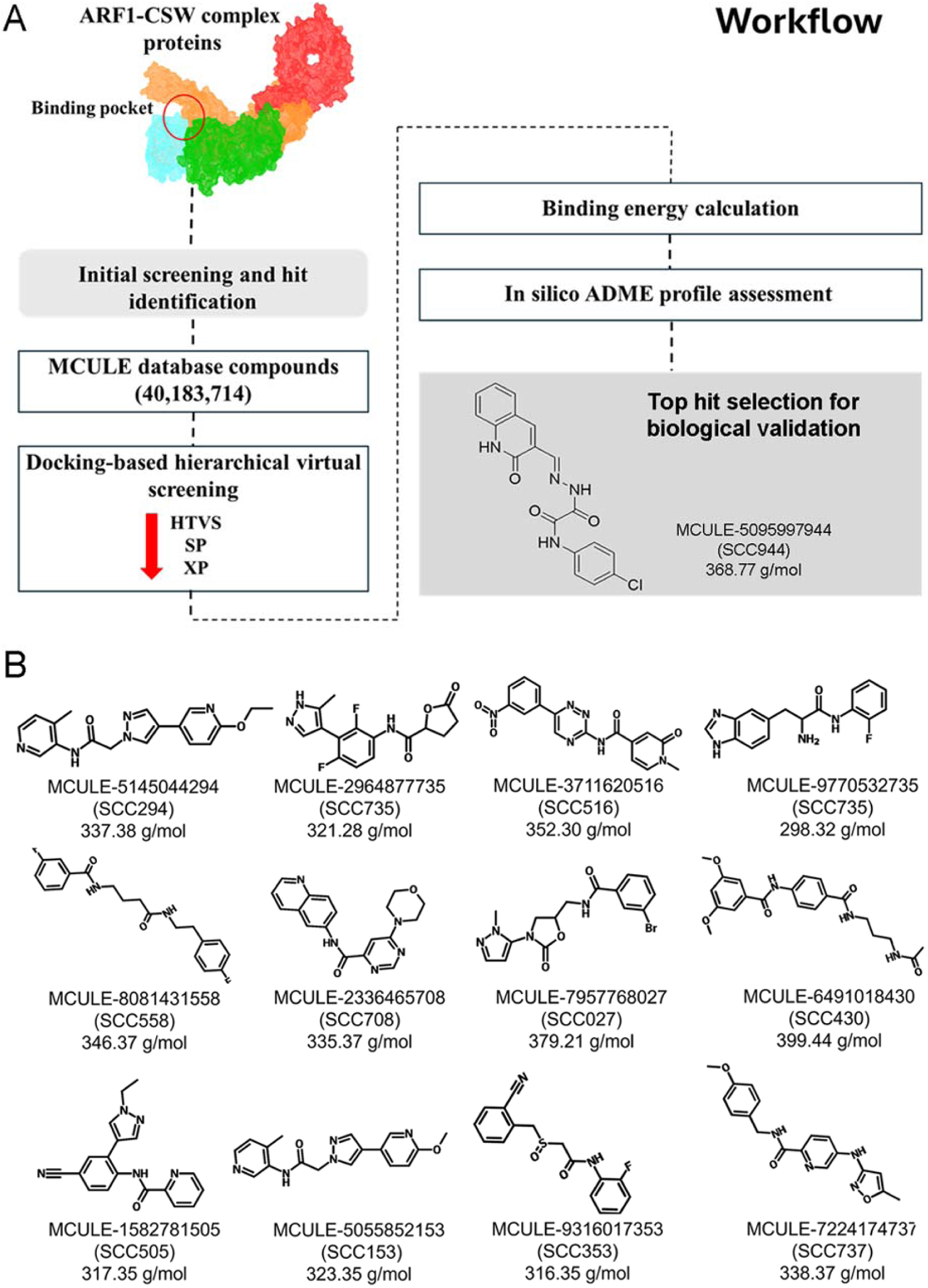
Structure based hierarchical virtual screening and identification of SMMs targeting the ARF1 interactions with CSW complex. **(A)** Schematic stepwise workflow for the identification of SMMs targeting ARF1-CSW complexes. SCC944 emerged as the top hits for further biological validation. **(B)** Additional top 12 hits that showed favorable binding scores and free energy ranking as well as strong SMM-CSW interaction pattens.

The superimposition of three representative compounds **MCULE-2336465708 (**Hereafter as **SCC708)**, **MCULE-5055852153** (Hereafter **SCC153**) and **MCULE-5095997944** (Hereafter **SCC944**) at the interface of the ARF1-CSW complex is shown in **Figure 4A**. These compounds exhibited interactions with various residues of ARF1, SMCR8 and C9orf72 *via* hydrogen bonds, halogen bonding and hydrophobic interactions (**Figures 4B-D, Supplementary Figure S1** and **Table S1).** For instance, the nitrogen atom of the quinoline ring and oxygen atom of the carbonyl group of **SCC944** formed π-π stacking and hydrogen bond interactions with His^65^ of C9orf72 and Ile^49^ of ARF1. Additionally, the nitrogen atom of the amide and chloro group of **SCC944** interacted with Pro^107^ of SMCR8 protein through hydrogen binding and halogen bond interactions, respectively (**Figure 4B** and **Figure S1**). **SCC708** exhibited hydrogen bond interactions with both SMCR8 and ARF1 residues Pro^107^ and Ile^49^ (**Figure 4C**). As for interactions observed for **SCC153**, the nitrogen atom of the amide and the nitrogen atom of the quinoline ring formed hydrogen bonds with Gly^109^ and Arg^151^ of SMCR8 respectively (**Figure 4D**). This analysis supported that these top compounds interact with ARF1, C9orf72, and SMCR8 within the ARF1-CSW complex.

**Figure 4.**
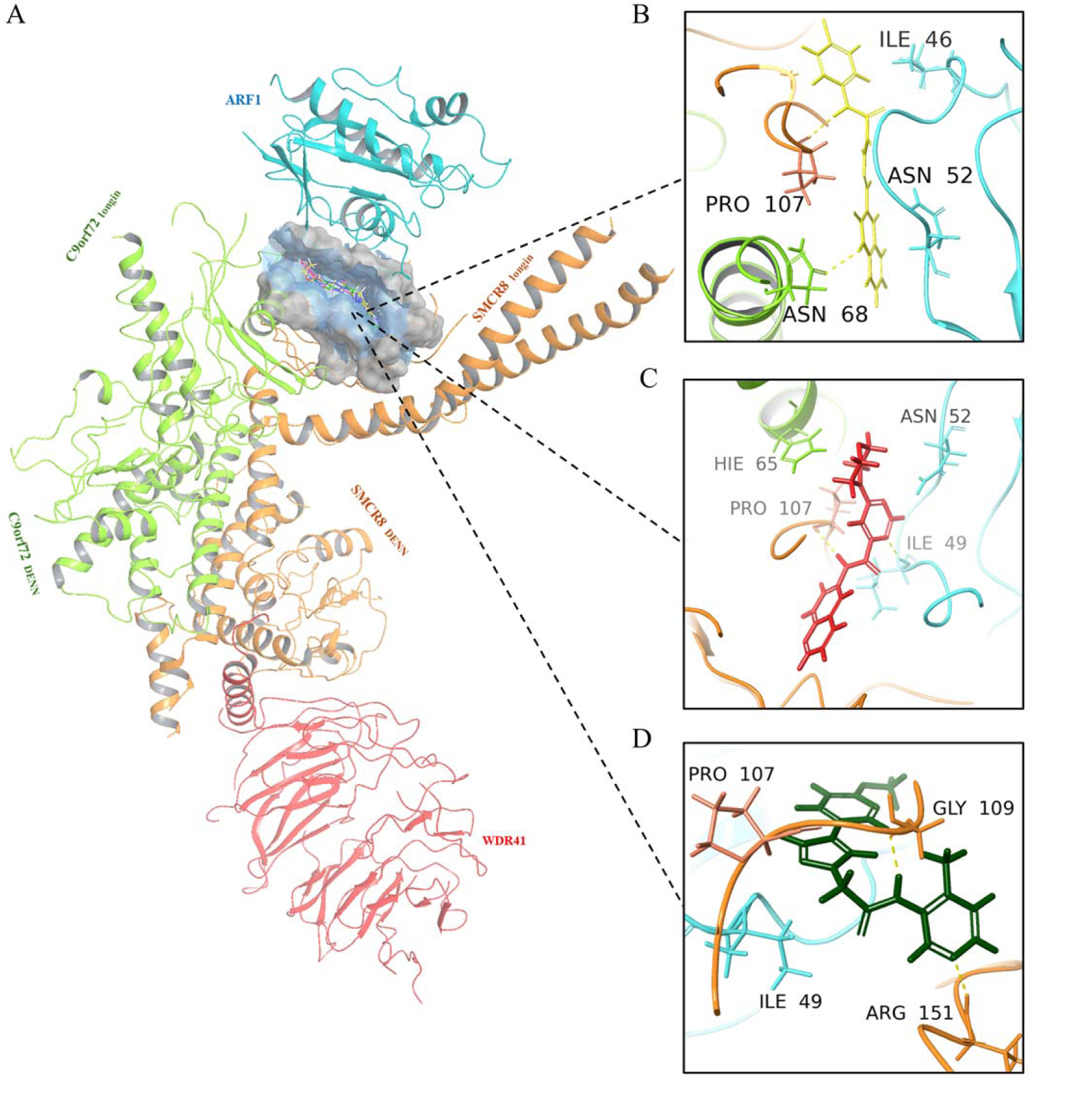
Binding modes of SMM targeting ARF1-CSW complex. (**A**) Superimpose of identified compounds **SCC944**, **SCC708** and **SCC153** into ARF1-CSW complex proteins (ARF1 protein is in transparent cyan, SMCR8 and C9orf72 proteins are in orange and green**, respectively** as well as WDR41 is shown in red color (PDB: 7MGE)). Identified compounds show different poses with a certain extent of overlap with each other. **(B-D)** 3D representation of identified compounds interactions with ARF1-CSW complex proteins. **SCC944** (yellow), **SCC708** (red) and **SCC153** (green) bind to interface of ARF1, C9orf72, SMCR8 *via* different interactions such as hydrogen bonds with Ile^49^, Gly^50^ of ARF1 and Pro^107^ of SMCR8, as well as π-π interaction with Hie^65^ of C9orf72.

Additionally, the ligand binding free energies of the top-ranking compounds were calculated using MM/GBSA analysis to predict the binding affinity to the ARF1-CSW protein complex. They ranged from -41.38 to -71.41 kcal/mol, indicating strong and stable interactions at the interface of ARF1 and CSW complex. Notably, **SCC944,** with a binding free energy of -60.89 kcal/mol, formed interactions with all three proteins, ARF1, SMCR8, and C9orf72, at the ARF1-CSW complex interface (**Supplementary Table S1)**.

Furthermore, the *in-silico* ADME/Tox analysis revealed that these top-ranking compounds exhibited favorable pharmacokinetics (PK) properties and the absorption, distribution, metabolism, excretion and toxicity (ADME/Tox) profiles (**Supplementary Table S2** and **S3)**. Using the ProTox-II server (18), the LD_50_ value for **SCC944** was predicted to be 3000 mg/kg, placing it in a low toxicity class (**Supplementary Table S4)**. Following this comprehensive workflow, **SCC944** emerged as a promising candidate, and was selected as the top compound for biochemical and cell-based biological validation.

### Direct binding of SCC944 to ARF1

To validate the *in-silico* prediction of **SCC944** interaction with ARF-CSW complex, we assessed the binding affinity and kinetics by surface plasma resonance (SPR). To determine the direct binding of the three representative compounds to ARF1, we covalently immobilized purified ARF1 to CM5 sensor chips and varied small molecule concentrations. The response was shown to increase with increased small molecule concentration and provided individual steady state affinity values (**Figure 5A-D**). This value was 167 nM for **SCC944**. As a comparison, SPR profiles showed 517 nM for **SCC708** and 249 nM for **SCC153 (Figure 5E)**. We also assessed the SPR interaction of ARF1 with a known non-binder, ZCL278, which selectively binds to Cdc42 small GTPase in the distinct Rho/Rac/Cdc42 subfamily (19, 20). We found a binding affinity of 1.41 mM for ZCL278 (**Figure 5D and E**), which confirms the strong binding of the small molecules **SCC944, SCC708 and SCC153** to ARF1.

**Figure 5.**
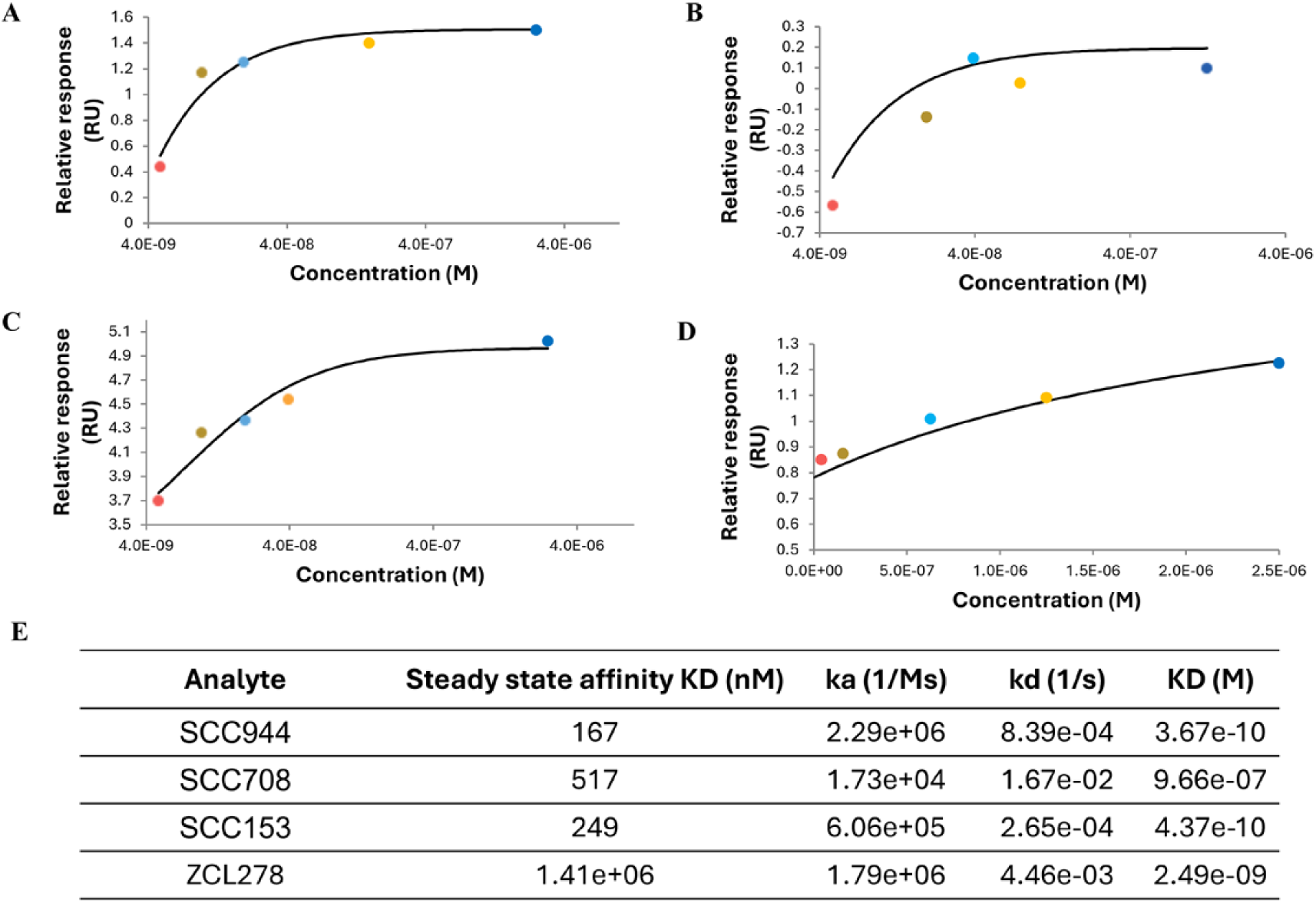
SMM-ARF1 binding evaluation by SPR. Binding response of **(A) SCC944**, **(B) SCC153**, **(C) SCC708** and **(D)** ZCL278 to the purified ARF1 protein using multi-cycle kinetics model and varying concentrations of 4.88 nm to 1.25 µM on Biacore 1S+ SPR. **(E)** Binding kinetics and affinity data for **SCC944**, **SCC153**, and **SCC708**.

### Effect of SCC944 on ARF1 activation and Golgi organization

To determine the biological effects of **SCC944** on ARF1 activation, HEK293 cells were treated with the compound and the basal GTP-bound ARF1 levels were determined (**Figure 6A, left panel**). BFA was employed as a well-known ARF inhibitor. The active GTP-bound ARF1 level was visibly reduced following treatment with BFA although the decrease was not statistically significant due to large errors (∼ 22% reduction; n=9). There was a more moderate decrease when treated with **SCC944** although not statistically significant (< 10%; n=9). Next, HEK293 cells were treated with 5’-Guanylyl imidodiphosphate (GDPNP), a non-hydrolysable analog of GTP, as a known ARF1 activator. When HEK293 cells were incubated with BFA and SCC944 followed by GDPNP treatment, both BFA and **SCC944** significantly suppressed the active GTP-bound ARF1 levels compared to GDPNP treatment alone (**Figure 6A, right panel**). These data indicated that both BFA and **SCC944** antagonized ARF1 activation.

**Figure 6.**
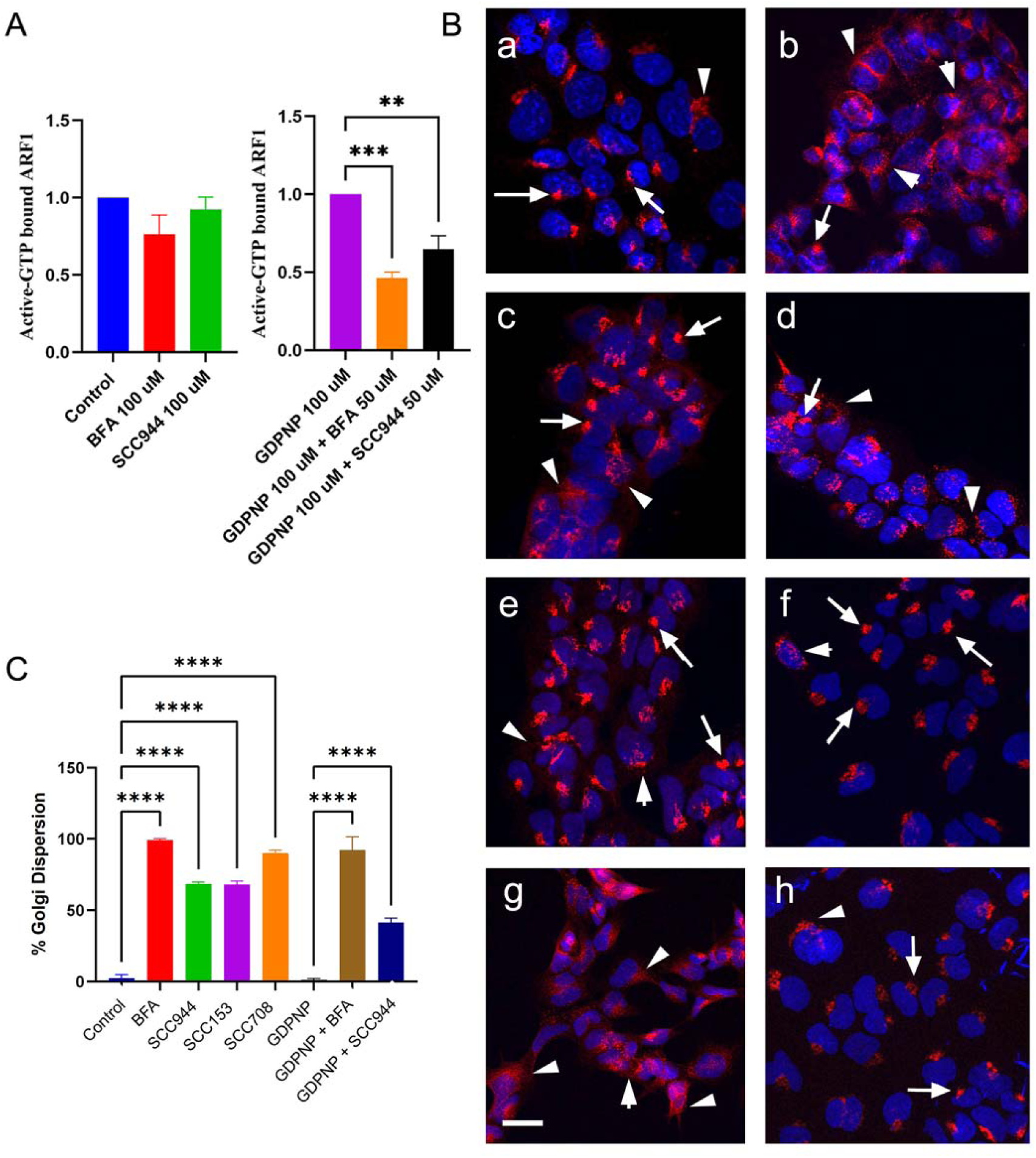
Biological functionality modulated by SMMs targeting ARF1-CSW interactions I - Golgi organization. **(A)** G-LISA analysis of ARF1-GTP levels in HEK293 cells treated with Control vehicle, 50 µM BFA, and 50 µM **SCC944** (Left panel) or with **100 µM** GDPNP, GDPNP+50 µM BFA, and GDPNP+ 50 µM **SCC944** (Right panel). All data are presented as mean ± SEM from duplicates from three independent experiments. ANOVA compared treatments to control (*P*-values ** *p*< 0.005, \*\*\**p* < 0.001, \*\*\*\**p* < 0.0001 were considered significant). **(B)** Effects of **SCC944, SCC708, SCC153**, GDPNP and BFA (50 µM) on the Golgi structures in HEK293 cells. Cells were fixed and stained with anti-GM130 antibody (Red). Cell nucleus was stained with DAPI (Blue). Immunofluorescent staining of GM130 in HEK293 cells treated with (a) Control vehicle, (b) BFA, (c) 50 µM **SCC944**, (d) 50 µM **SCC153**, (e) 50 µM **SCC708,** (f) GDPNP, (g) GDPNP followed by BFA, (h) GDPNP followed by 50 µM **SCC944**. Arrows point to the perinuclear compact Golgi. Arrowheads point to dispersed Golgi in cytoplasm. Bar: 30 µm. **(C)** Bar graph showing percent dispersion of Golgi apparatus with and without the treatments. The mean percent Golgi apparatus dispersion was compared. All data are presented as mean ± SEM from duplicates from three independent experiments. ANOVA compared treatments to their respective control (*P*-value \*\*\*\**p* < 0.0001 was considered significant).

Golgi apparatus as a steady-state structural organization depends on ARF1 (21, 22). To establish whether the SCC944 modulation of ARF1 activation impacted on functional cellular phenotypes, we determined the effects of **SCC944, SCC708, SCC153**, GDPNP and BFA on the Golgi organization in HEK293 cells (**Figure 6B: a-h**). In most control cells, the cis/medial Golgi marker GM130 exhibited a distinct compact perinuclear localization (**Figure 6B: a, arrows**). Treatment with BFA resulted in not only fragmentation but also near-complete dissolution of the Golgi structure, with GM130 immunoreactivity becoming widely dispersed throughout the cytoplasm (**Figure 6B: b, arrowheads**). Golgi apparatus in cells treated with 50 µM **SCC944** and **SCC153** scattered to some extent (**Figure 6B: c and d, arrowheads**) but led to approximately 70% Golgi dispersal (**Figure 6C**), suggesting a less disruptive effect on Golgi integrity. The scattering of Golgi apparatus in cells treated with **SCC708** was more widespread than that of **SCC944** and **SCC153** treatment (**Figure 6B: e, arrowheads**) and showed approximately 90% Golgi dispersion closer to the BFA effect (**Figure 6C)**. GDPNP treated cells showed perinuclear Golgi localization although the Golgi structures were not as compact as in control (**Figure 6B: f, arrows**). Cells treated with GDPNP followed by BFA showed extensive Golgi dispersion similar to that from BFA treatment alone (**Figure 6B: g, arrowheads).** Some Golgi structures in the cells treated with 50 µM **SCC944** in the presence of GDPNP were not as compact as those in the control or cells treated with GDPNP alone (**Figure 6B: compare a, f, and h, arrowheads**). Nevertheless, the ∼ 40% Golgi dispersion observed with SCC944 (**Figure 6C**) is consistent with ARF1 modulation that perturbs, but does not severely dismantle, Golgi organization, in contrast to BFA treatment (**Figure 6B, h**, and **Figure 6C**).

### Effect of SCC944 on mitochondrial and microtubule organization

Studies in yeast indicated that besides the canonical roles of ARF1 in Golgi organization and vesicular trafficking, ARF1 is involved in mitochondrial dynamics and homeostasis (23, 24). In HEK293 cells, mitochondria positive for anti-citrate synthase (CS) immunoreactivity showed a widespread cytoplasmic distribution of elongated interconnected tubule structures, small fragmented or circular morphology, or intermediate feature with shorter rod-like structures with less connections as previously analyzed (**Figure 7A, Control; green and merged panels; Figure 7B**), (25). BFA treatments, however, resulted in the tendency of withdrawal of mitochondrial from cell periphery towards the perinuclear region with fragmentation shown as small circular morphology (**Figure 7A and B, BFA**). On the other hand, HEK293 cells treated with 50 µM **SCC944** showed less complete fragmentation of mitochondria (**Figure 7A and B, SCC944**). When the HEK293 cells were stimulated with GDPNP, the mitochondrial distribution resembled control with low fragmentation (**Figure 7A and B, GDPNP**), while cells treated with 50 uM BFA and then stimulated with GDPNP, showed lower fragmentation than BFA alone (**Figure 7A and B, GDPNP+BFA**). **SCC944** stimulated with GDPNP treatment showed the most prominent change from being slightly fragmented to almost 75% mitochondrial fragmentation (**Figure 7A and B, GDPNP+SCC944**), supporting that ARF1 plays important roles in regulating mitochondrial distribution and functions.

**Figure 7.**
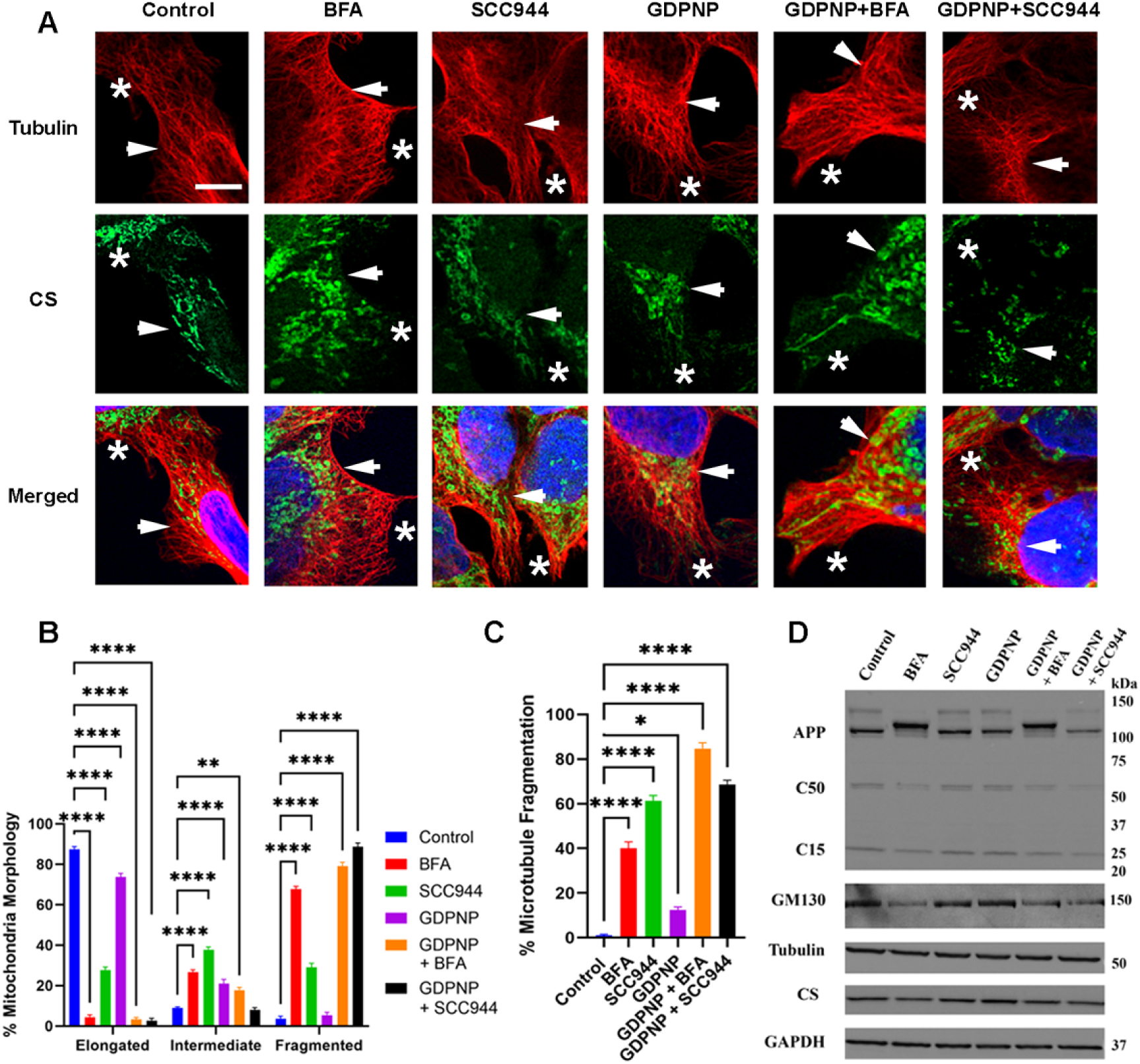
Biological functionality modulated by SMMs targeting ARF1-CSW interactions II – Mitochondrial and microtubule organization. **(A)** Effects of BFA, GDPNP, and **SCC944** on the mitochondrial and microtubule distribution in HEK293 cells. Cells were fixed and then stained with anti-CS (Green) or anti-tubulin (DM1α) (Red). Cell nucleus was stained with DAPI (Blue) and immunofluorescent stain on HEK293 cells treated with control vehicle, BFA, **SCC944**, GDPNP, BFA followed by GDPNP, and **SCC944** followed by GDPNP. Arrows point to distribution in the perinuclear region and asterisks point to the cytoplasmic region. Bar: 30 µm. **(B)** Percent mitochondrial alterations with and without treatments. All cells were given an elongated, intermediate, or fragmented phenotype and the percentage of each was compared between treatments. All data represented as mean ± SEM from duplicates from three independent experiments. ANOVA compared treatments to their respective control (*P-*values \*\**p<*0.01 and \*\*\*\**p<*0.0001 were considered significant). **(C)** Percent microtubule fragmentation with and without treatments. All cells were given a score of microtubule fragmentation between 0 (long, continuous, thread-like) to 100 (completely fragmented) and the percentages were compared between treatments. All data represented as mean ± SEM from duplicates from three independent experiments. (*P-*values \**p<*0.05 and \*\*\*\**p<*0.0001 were considered significant). **(D)** Representative western blots from control and treated HEK293 cell lysates. All treatments were analyzed for expression levels in APP, GM130, α-tubulin, citrate synthase (CS), and GAPDH. The molecular weights were indicated on the right side.

Mitochondrial integrity is often associated with microtubule stability. To compare microtubule fragmentation, we used a scoring system previously described on axons to manually create a microtubule fragmentation index in HEK293 cells (26). We observed a well-developed microtubule network in HEK293 cells with features of long, continuous and thread-like microtubules (**Figure 7A, Control; red panels, asterisk).** In BFA treated cells, microtubules are shorter and disorganized near the cell periphery (**Figure 7A, BFA**). Similar disorganized microtubules can be seen in **SCC944** treated cells (**Figure 7A, SCC944; red panels, arrow**). HEK293 cells stimulated with GDPNP showed similar well-developed microtubule network as untreated cells although with more fragmentation around the perinuclear region (**Figure 7A, GDPNP**). BFA and **SCC944** treatment after GDPNP stimulation showed more overall fragmentation and disorganized microtubules on the cell periphery than the treatments alone (**Figure 7A and C**).

### Effect of SCC944 on protein expression and post-translational processing

Along with the regulation of endoplasmic reticulum (ER)-Golgi organization, ARF1 dysregulation can impair protein processing and sorting through ER-Golgi structures. Compared to the control cells, the cis/medial Golgi resident GM130 expression was reduced when HEK293 cells were treated with BFA (**Figure 7D, GM130**). Both **SCC944** and GDPNP treated cells showed similar GM130 expression (**Figure 7D, GM130**). GM130 expression level was not recovered in the cells treated with BFA or **SCC944** with GDPNP stimulation (**Figure 7D, GM130**).

Amyloid precursor protein (APP) undergoes post-translational maturation through ER-Golgi and produced carboxyl-terminal fragments (CTFs) in the vesicular trafficking network, which have the potential to aggregate into Aβ (27, 28). HEK293 cells express holo APP as a doublet with immature upper band and a mature lower band at ∼ 100 kDa (29, 30). We also detected ∼ 55 kDa and ∼ 25 kDa CTFs (**Figure 7D, APP, C50 and C15**)(31). As expected, BFA treatment suppressed APP maturation as well as reduced CTFs, which were not recovered with GDPNP treatment (**Figure 7D, APP, C50 and C15**). **SCC944** and GDPNP did not change the pattern of APP expression and processing. However, when HEK293 cells treated with **SCC944** in the presence of GDPNP, the APP total expression and proteolytic processing were reduced, and its proteolytic pattern by **SCC944** is different from BFA treatment, indicating that **SCC944** and BFA affected APP processing differently.

## Discussion

Previous studies on ARF1 modulators included BFA and LM11 which inhibit ARF1 activation by GEFs (32–34) and protect motor neurons from degeneration (35, 36). However, considering the ALS/FTD relevant GAP activity of CSW complex for ARF1 (9, 17), our study represented an important step towards development of pathophysiological relevant therapeutic SMMs that directly target the ARF1-CSW interface.

We hypothesized that the loss of function mutations in *C9orf72* leads to local accumulation of ARF1-GTP at sites of CSW activity, and the elevated levels of ARF1-GTP play an important role in the pathogenesis of ALS/FTD. This hypothesis is supported by our findings that ARF-GAP protein ASAP1 is inactivated in the posterior frontal motor cortex of both sporadic ALS and ALS patients with *C9orf72* mutations.

It is interesting to note that ASAP1 activity is reduced in both sALS and C9ALS patients. Since ARF1 activity would have already been upregulated due to *C9orf72* haploinsufficiency in C9ALS, one plausible explanation for downregulating ASAP1 in C9ALS would seem to be a mechanism to ensure ARF1 hyperactivity in ALS/FTD. This notion can also be implied in the C9orf72 ectopic expression studies. Introducing C9orf72 into HEK293 cells resulted in the Golgi dispersion like the BFA treatment that inhibits ARF1. These studies suggested that in C9ALS, where ARF1 likely already remains in an activated GTP-bound form due to loss-of-function *C9orf72* mutations, ASAP1 is also inactivated to keep ARF1 hyperactivated. Therefore, ARF1 activity may be beneficial for brain function but its overactivation can put vesicle trafficking on overdrive and are thus damaging to cellular homeostasis. Therefore, our data not only supports ARF1 overactivation as a pathogenic event in ALS, but they are also consistent with the literature that ARF1 is important for embryonic development (37).

Through structure-based hierarchical virtual screening, **SCC944** emerged as a promising modulator with strong binding affinity and favorable ADME/Tox properties. This compound interacts with various residues in ARF1, SMCR8, and C9orf72 proteins. Notably, it interacts with key residues essential for GAP activity, including Gly^50^ and Ile^49^ of ARF1, where mutations have been shown to diminish or abolish GAP function (38). Given that the binding pocket was determined at the interface of ARF1, C9orf72 and SMCR8, it is reasonable that the initial hit compounds would not interact with the WDR41 protein in ARF1-CSW complex. Indeed, previous studies suggested that WDR41 in the CSW complex is not essential to the GAP activity for specific small GTPase (10, 11).

Both **SCC944** and BFA caused varying levels of Golgi dispersion, mitochondrial disorganization as well as microtubule fragmentation, consistent with their being ARF1 modulators. However, **SCC944** and BFA affected Golgi-mediated APP maturation and proteolytic processing differently. Based on these observations, we propose a working model illustrating that BFA and SCC944 may regulate location biased ARF1 functions through distinct mechanisms (**Figure 8**).

**Figure 8.**
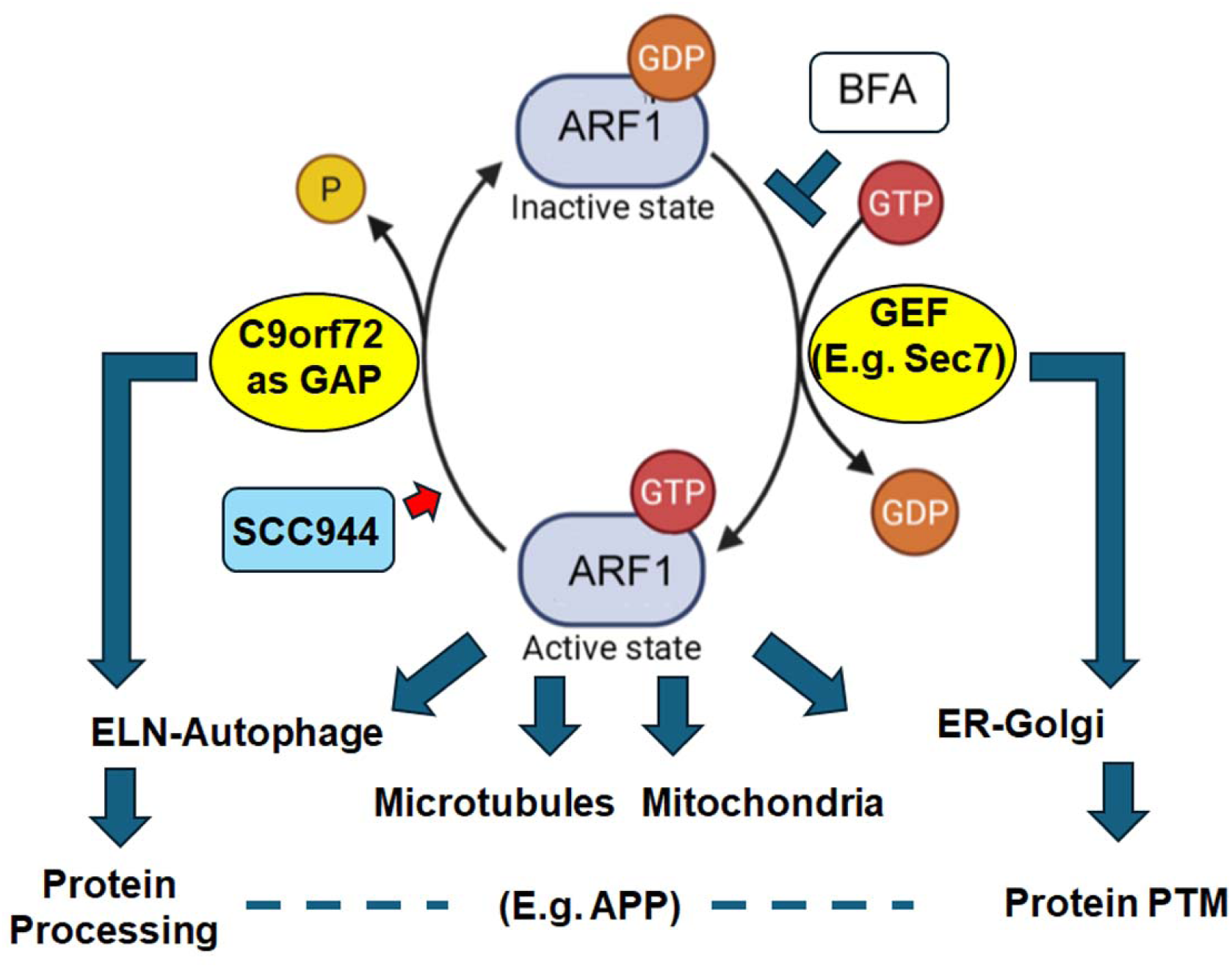
Schematic model on differential regulation of intracellular organelle organization by ARF1 and SCC944. ELN: Endolysosomal Network; PTM: Posttranslational Modifications.

ARF small GTPases regulate both the cellular organelle structural integrity and protein trafficking. The molecular mechanism of our observed APP maturation and processing may break down into several components. First, as expected, BFA inhibited ARF1 activation and reduced the cis/medial-Golgi matrix protein GM130. Because GM130 acts as a structural anchor that stabilizes Golgi stacks and coordinates glycosylation enzymes, Golgi structures lacking GM130 would fragment or collapse into ER. Therefore, APP cannot complete normal maturation with glycosylation, leaving it as an intermediate form before reaching complete maturation. On the other hand, **SCC944** did not clearly suppress the basal ARF1 activity and kept normal GM130 expression, likely preserving the baseline structural integrity of the Golgi. Consequently, both the cellular expression of APP and its standard glycosylation pathways remain intact. Then, when ARF1 is hyperactivated by GDPNP, SCC944 acts as a competitive antagonist. It displaced the full endogenous activator and suppressed ARF1 activity towards the basal level. This sudden drop in ARF1 signaling may be severe enough to downregulate GM130 as shown in **Figure 7D**.

This model proposes that when **SCC944** produced a more moderate ARF1 inhibition (**Figure 6**) and selectively impaired vesicle trafficking without fully destroying the internal enzymatic environment of the Golgi like that by BFA, newly synthesized APP trapped in the ER-Golgi compartments can trigger a quality-control mechanism such as ER-associated degradation (ERAD) or autophagy. In this case, the trapped APP can get degraded by cytoplasmic ubiquitin or proteasome system. Meanwhile, APP remained in the Golgi is still glycosylated whereas APP glycosylation was severely affected by BFA.

This model is supported by the literature that (1) BFA interacts with the switch II domain of ARF1-GDP and the hydrophobic groove of Sec 7 domain and (2) Sec7 GEF is localized to Golgi to support Golgi maturation (39–41). This mechanism can explain the strong impact of BFA on APP processing whereas **SCC944** did not diminish APP maturation. On the other hand, **SCC944** demonstrated greater impact on mitochondria fragmentation (**Figure 7D**). Given that ARF1 is also localized within mitochondria (23) and regulates fatty acid metabolism (24), our findings highlight the pleiotropic roles of ARF1 and the potential pathophysiological importance of targeting mitochondrial ARF1. Mitochondrial dyshomeostasis represents a critical and early pathogenic mechanism in motor neuron degeneration associated with ALS/FTD (42).

Finally, mitochondrial redistribution and microtubule fragmentation when stimulated with GDPNP increase when combined with the BFA treatment. **SCC944** showed a similar impact on both mitochondria and microtubule disorganization with and without GDPNP present. On the other hand, BFA showed more perinuclear mitochondria and fragmented microtubules, only when combined with GDPNP. This is consistent with the literature as shown that BFA when combined with an ARF1 GEF, locks the pathway and can overactivate other small GTPases that would catastrophically damage microtubule remodeling as well as the mitochondrial redistribution (43). **SCC944** does not show the same phenotype, suggesting that the inhibition of ARF1-CSW is regulated in different pathways. Accordingly, modulation of ARF-GAP functions within the ARF1-CSW complex may offer a selective and effective therapeutic strategy for ALS/FTD.

## Materials and Methods

### Docking based hierarchical virtual screening

#### Protein preparation

The crystal structure of C9orf72: SMCR8:WDR41 (CSW) in complex with ARF1 was obtained from the Protein Data Bank (https://www.rcsb.org/structure/7mge) using the PDB entry code 7MGE. To prepare the proteins, the protein preparation wizard module of the Schrödinger software package was employed (44). The co-crystallized GDP was deleted, hydrogen atoms were added, and water molecules were removed. Then, the OPLS4 force field was used for optimization and minimization of the ligand structures.

#### Grid generation

The site Map tool from Schrödinger (45) was utilized to identify the binding pockets of the ARF1-CSW in the complex. The highest-ranking binding pockets were located around the co-crystallized GDP molecule. A grid generation was generated around the GDP binding pocket at the interface of CSW and ARF1, encompassing the following residues: Ile^49^, Gly^50^, Phe^51^, Thr^48^ of ARF1 and Pro^108^, Ser^110^, His^106^, Val^150^, Arg^147^, Pro^152^, Arg^151^ of SMCR8 as well as Phe^61^, His^65^, Asn^64^ and Asn^68^ of C9orf72. The assigned coordination values were as follows: X=133.56, Y=137.7 and Z=165.1.

#### Ligand preparation

The MCULE database, which contains 40,183,714 chemicals, was sourced from the MCULE library (https://go.drugbank.com/). The top 3,000 identified small molecules derived from the virtual screening on the MCULE web server were utilized as a screened library for local docking-based hierarchical screening. The LigPrep module of Schrödinger software was employed to prepare, neutralize, desalt, and adjust the tautomer’s of the compounds, followed by energy minimization using the OPLS4 force field (46). LigPrep, Schrödinger, LLC, New York, NY, 2023).

#### Structure-based virtual screening and hierarchical molecular docking

Initially, a total of 40,183,714 chemicals were screened against the identified binding pocket at the interface of the CSW complex with ARF1 proteins using the open-source MCULE web server. This initial docking-based screening yielded 3,000 compounds with the highest docking scores. Subsequently, docking studies were performed using the Glide module of Schrödinger software. For the docking-based hierarchical virtual screening, the top-ranked compounds were docked at three levels of accuracy: high-throughput virtual screening (HTVS), standard precision (SP), and extra precision (XP). The screening criteria were established to retrieve 10%, 20%, and 20% of compounds after HTVS, SP, and XP docking, respectively (47).

#### Binding free energy calculation with MM/GBSA

The 14 top-ranked compounds identified from the hierarchical virtual screening were selected for <u>m</u>olecular <u>m</u>echanics energies combined with <u>g</u>eneralized <u>B</u>orn and <u>s</u>urface <u>a</u>rea continuum solvation (MM/GBSA) analyses. MM/GBSA was utilized to approximate the binding free energy of small molecules (ligands) to biological targets (48). To evaluate the free energy, the Prime MM-GBSA module of the Schrödinger Suite was employed. The ligand binding free energy is determined by the following equation:

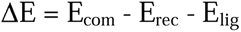

Where ΔE, E_com_, E_rec_, and E_lig_ represent the free binding energy, the energy of complex, receptor and ligand energy, respectively. The binding free energies of compounds were calculated with VSGB implicit solvation and OPLS3 force field, and the residues within 10Å distance from the ligand were kept flexible during the analysis.

#### *In silico* ADME/Tox profile assessment

The physicochemical properties, pharmacokinetics, and drug compatibility of the top-ranked compounds identified from the hierarchical virtual screening, as well as the novel designed compounds, were predicted using the QikProp tool from Schrödinger (QikProp, Schrödinger, LLC, New York, NY, 2024). The ADME/Tox properties relevant to the drug development process—including absorption (water solubility, human oral absorption), distribution (blood-brain barrier permeability, total solvent-accessible surface area, and prediction of binding to human serum albumin), metabolism, and excretion (predicted apparent MDCK and Caco-2 cell permeability)—were also evaluated. To assess the safety of the top-ranked compounds derived from the hierarchical virtual screening and the novel modified compounds, we utilized the QikProp tool from Schrödinger and the ProTox-II server (18) for an initial toxicity evaluation. This assessment provided insights into various toxicity profiles, including hepatotoxicity, carcinogenicity, immunotoxicity, mutagenicity, and cytotoxicity of the compounds.

#### Scrutinization of potential ARF1-CSW complex modulators

A thorough evaluation process was conducted to identify a hit compound with favorable binding characteristics and optimal ADME/Tox properties for the generation of an optimized library. Among the top-ranked compounds obtained from the hierarchical virtual screening, **SCC944** was selected as the seed compound for further optimization due to its promising interaction profile, favorable binding energy, and good ADME/Tox profile.

#### Bioinformatic analysis

The protein-protein interactions of differentially expressed proteins were analyzed by STRING (http://string.embl.de) with the following analysis parameters: Species-Homo Sapiens, meaning of network edges-evidence, active interaction sources-all, interaction score-0.150.

### Biological assays

#### Ligand-ARF1 binding by SPR

Surface plasmon resonance (SPR) experiments were performed on a Biacore 1S+ using CM5 sensor chip (Cytiva, Marlborough, MA). All solutions used in the experiment were prepared with Mill-Q water, filtered with a 0.22-μm membrane filter, and degassed before use. Recombinant human ARF1 protein (Creative Biomart Inc, Shirley, NY) in Tris solution (20 mM, pH 7.4) at a concentration of 1 mg/mL was diluted to 50 μg/mL with sodium acetate buffer (pH 4.0). The chip was activated with EDC/NHS (20 μL/min for 100 s). Then, ARF1 was loaded (20 μL/min for 600 s) and immobilized covalently. Approximately 10,000 RU of ARF1 was immobilized on the chip. Any excess of unbound ARF1 was removed by flowing Tris solution [20 mM, pH 7.4, with 5% (wt/vol) DMSO]. ZCL278, **SCC944**, **SCC708** and **SCC153** were prepared as 4.88-nm to 1.25-μM solutions in Tris solution [20 mM, pH 7.4, with 5% (wt/vol) DMSO], and injected (10 μL/min for 100 s). After each loading, data was collected and analyzed with the Biacore Insight Evaluation software.

#### Reagents and cell lines

G-LISA kit (BK132) to measure ARF1 activity were purchased from Cytoskeleton, Inc. (Denver, CO). **SCC944**, Brefeldin A and GDPNP were purchased from Mcule (CA, USA), Millipore Sigma (MD, USA) and Santa Cruz Biotechnology (Texas, USA), respectively. HEK 293 cells were purchased from American type Culture Collection (ATCC) (Manassas, VA) and cultured as described. Compounds were administered to cells in 0.5% DMSO (Sigma Aldrich) with further dilutions in culture medium. Antibodies used for western blot and immunofluorescence in this study include: Mouse anti-GM130 (610822, 1:500 for WBs and 1:100 for IF, BD Transduction Laboratories), Rabbit anti-β amyloid precursor protein (CT695) (512700, 1:2000 for WB, Invitrogen), Rabbit anti-Citrate synthase (326-340) (C5498, 1:500 for WB and 1:100 for IF, Sigma Aldrich), Mouse anti-α-tubulin (T9026, 1:500 for WB and IF, Sigma Aldrich), Mouse anti-GAPDH (CB1001, 1:1000 for WB, Calbiochem). For WB, primary antibodies were detected using Anti-Mouse IgG, HRP conjugate or Anti-Rabbit IgG, HRP conjugate (1:2500, Promega). For IF, primary antibodies were detected using anti-rabbit Alexa Fluor 488 (A11008, 1:600, Invitrogen) or Cy3 conjugated anti-mouse IgG (715165150, 1:600, Jackson Immunoresearch).

#### Cell Culture and Transfection

The HEK 293 kidney epithelial cells were obtained from ATCC (CRL-1573, ATCC, Manassas, VA). They were grown at 37°C and 5% CO² on glass coverslips in Eagle’s minimum essential medium (ATCC) containing 10% fetal bovine serum, and 1% Penicillin-Streptomycin (Invitrogen). When the cells reached around 70% confluency, lipofectamine 2000, *pEGFP-C9orf72* (49) and serum free medium were added to cells to let grow for 4 hours at 37°C. After 4 hours, half of the medium was removed and replaced with medium containing serum. After 18 hours, all medium was removed and replaced with fresh serum medium and let grow for 24 hours. Cells were fixed on coverslips with paraformaldehyde for 20 minutes.

#### Cell treatment and immunofluorescence light microscopy

The HEK 293 cells were grown at 37°C and 5% CO² on glass coverslips in Eagle’s minimum essential medium (ATCC) containing 10% fetal bovine serum, and 1% Penicillin-Streptomycin (Invitrogen). When the cells reached around 70% confluency, they were treated with **SCC944, SCC708, SCC153**, GDPNP or BFA (50 µM and 100 µM) for 1.5 hrs, followed by a 30-minute treatment of GDPNP to some BFA and **SCC944** coverslips. After treatment, cells were fixed with paraformaldehyde for 20 min. Cells were washed with PBS (3X), and blocked using BSA (5%) in PBS followed by incubation at room temperature for 1 hr. Next, cells were washed with PBS and incubation with primary antibody was performed in phosphate-buffered saline containing 2.5% bovine serum overnight at 4 °C. After phosphate buffered saline washes, staining with the secondary antibody was carried out for 60 min at room temperature. Images were collected with a Zeiss Axio Observer 7 inverted microscope (Carol Zeiss, Thornwood, NY).

#### Western Blot

Treated cells were lysed using Pierce RIPA Buffer (Thermo Scientific) and protein concentrations were measured using the Pierce BCA Protein Assay Kit (Thermo Scientific). All lysates were diluted to 15 ug and loaded into an 8%-16% gradient Tris-glycine Gel (Invitrogen). The protein was transferred onto a 0.2 µM nitrocellulose membrane and blocked for 1 hour in 5% nonfat dry milk in TBST (Tris-buffered saline + 0.1% Tween 20). After blocking, primary antibody was added overnight. The membrane was then briefly washed, and the corresponding HRP-conjugated secondary antibody was added for 1 hour. The membrane was soaked in chemiluminescent substrate (SuperSignal Substrate, Thermo Scientific) for 5 minutes and imaged on ChemiDoc MP (BioRad).

#### G-LISA assay

HEK293 (75% confluent) were treated with different concentrations (50 and 100 µM) of **SCC944** and BFA compounds for 2 hours. For stimulated ARF1 G-LISA, GDPNP as ARF1 activator (100 µM) was used to activate ARF1 in HEK 293 cells for 30 minutes after treatment of **SCC944** or BFA for 1 hour and 30 minutes. The cells were lysed 2 hours later and ∼ 1mg/ml protein was used for G-LISA analysis per manufacturer’s (Cytoskeleton) instructions. After placing the collected lysates in a 96-well ARF1-GTP binding plate, any unbound GTP was washed out and the bound GTP levels were measured on Biotek Synergy HT (Agilent, Winooski, VT).

#### Human brain tissues and immunohistochemistry

Immunohistochemistry (IHC) staining was done on 6 µm sections. Slides containing de-identified human brain tissues from the posterior frontal motor cortex, including normal control brains, sALS brains, and C9orf72 ALS brains from VA Biomedical Laboratory Research & Development Service (VABBB, Boston, MA), were used for staining (n=9). The slides were paraffin fixed and were deparaffinized in xylene and were rehydrated using decreasing concentration of alcohol. Antigen retrieval was then performed, and the slides were incubated with rabbit antigen-blocking buffer for an hour. These slides were then incubated with polyclonal Phospho-ASAP1 (Tyr782) antibody (Thermo Fisher Scientific, Waltham, MA)(1:100) for overnight at 4°C. After rinsing with PBS, Vectastain ABC reagent was added for an hour, followed by incubation with biotinylated secondary antibody (anti-rabbit) for 2 hours. The tissues were then exposed to DAB staining and mounted on coverslips. DAB staining images were acquired from Zeiss M-1 Axio Imager Fluorescence Microscope with AxioCam camera and 3-D software (Carol Zeiss, Thornwood, NY).

#### Statistical analysis

For all analyses, data were presented as mean±SEM from two to nine independent experiments with each having multiple replicates. Differences between treated groups and control were analyzed using one-way ANOVA. *P*-values \**p* < 0.05, \*\**p* < 0.01, \*\*\**p*< 0.001, \*\*\*\**p* < 0.0001 were considered significant. For IF staining analyses, phenotypic changes were scored and percentages were calculated by three different individuals using three independent experiments. For microtubule quantification, we used a scoring system previously described on axons to manually create a microtubule fragmentation index in HEK293 cells (26). Thus, 0% was shown as long, continuous, thread-like network, 33% represents partial fragmentation around the perinuclear regions with thread-like networks reaching into the periphery, 66% shows fragmented microtubules around the perinuclear regions with noticeable breakage around the cell edges, and 100% as complete loss of long filaments with diffused fragments into the cytoplasm.

## Supplementary Information

The online version contains supplementary material available.

Figs. S1

Tables S1 to S4

## Acknowledgements

We thank Shayan Nik Akhtar for technical assistance and VA Biomedical Laboratory Research & Development Service for ALS human biomaterials. This study is supported in part by NIH Director’s Transformative Research Award R01GM146257 for Accelerating Leading-edge Science in ALS (ALS^2^) Initiative (QL) and Smart State Center for Economic Excellence of South Carolina (QL).

## Authors’ contributions

ED-CADD, SPR, cell culture, transfection, Western blot, IFC, data analysis and manuscript writing. FA-CADD, G-LISA, IFC, cell culture, data analysis and manuscript writing. AJC-IFC, IHC, and data analysis. RT-cell culture, IFC, and imaging analyses. CB-G-LISA and data analysis. YHC and QL-concept and study design, data analyses and interpretation, and manuscript writing and editing. All authors read and approved of the final manuscript.

## Data Availability

All data supporting the findings of this study are available in the main text and/or the supplementary materials. The identities of some compounds docked in this study and the docking results, including number, SMILES, and docking score, are available upon request from the corresponding author in accordance with institutional technology safety and privacy policy.

## Declarations

## Ethical approval and Consent to participate

Not applicable per Institutional Research Board (IRB) committee guidance. The de-identified fixed human tissues were provided by VA Biomedical Laboratory Research & Development Service (VABBB).

## Consent for publication

All authors have approved of the consents of this manuscript and provided consent for publication.

## Abbreviations

AD: Alzheimer’s Disease
ADME/T: Absorption, Distribution, Metabolism, Excretion, and Toxicity ox
ALS: Amyotrophic Lateral Sclerosis
ARF1: ADP-ribosylation factor 1
BFA: Brefeldin A
C9orf72 CS: Chromosome 9 Open Reading Frame 72 Citrate Synthase
CSW: C9orf72-SMCR8-WDR41 Complex
FTD: Frontotemporal Dementia
GAP: GTPase-activating Protein
G-LISA: GTPase-linked Immunosorbent Assay
GDPNP: 5’-Guanylyl imidodiphosphate
MD: Molecular Dynamics
MM/GBS: Molecular Mechanics/Generalized Born Surface Area
PK: Pharmacokinetics
RAB: Ras-related in Brain
SMM: Small Molecule Modulator
SMCR8: Smith-Magenis Syndrome Chromosomal Region Candidate Gene 8
WDR41: WD Repeat Domain 41

## References

1. Mehta, P. et al. Recruitment of patients with amyotrophic lateral sclerosis for clinical trials and epidemiological studies: descriptive study of the National ALS Registry’s research notification mechanism. J. Med. Internet Res. 23, 28021; 10.2196/28021 (2021).

2. Priyadarshini, S. & Ajroud-Driss, S. Update on ALS treatment. Curr. Treat. Options Neurol. 25, 199–212; 10.1007/s11940-023-00757-4 (2023).

3. Gillingham, A. K. & Munro, S. The small G proteins of the Arf family and their regulators. Annu. Rev. Cell Dev. Biol. 23, 579–611; 10.1146/annurev.cellbio.23.090506.123209 (2007).

4. Renton, A. E., et al. State of play in amyotrophic lateral sclerosis genetics. Nat. Neurosci. 17, 17–23; 10.1038/nn.3584 (2014).

5. Majounie, E. et al. Frequency of the C9orf72 hexanucleotide repeat expansion in patients with amyotrophic lateral sclerosis and frontotemporal dementia: a cross-sectional study. Lancet Neurol. 11, 323–330; 10.1016/S1474-4422(12)70043-1 (2012).

6. Nik Akhtar, S., et al. Crosstalk between the Rho and Rab family of small GTPases in neurodegenerative disorders. Front. Cell. Neurosci. 17, 1084769; 10.3389/fncel.2023.1084769 (2023).

7. Adarska, P. et al. ARF GTPases and their ubiquitous role in intracellular trafficking beyond the Golgi. Front. Cell Dev. Biol. 9, 679046; 10.3389/fcell.2021.679046 (2021).

8. Arrazola Sastre, A., et al. Small GTPases of the Rab and Arf families: key regulators of intracellular trafficking in neurodegeneration. Int. J. Mol. Sci. 22, 4425; 10.3390/ijms22094425 (2021).

9. Su, Y. et al. Multi-omics responses in post-acute sequelae of COVID-19. Nature 603, 138–144; 10.1038/s41586-022-04569-9 (2022).

10. Nörpel, J. et al. Structure of the human C9orf72–SMCR8 complex reveals a multivalent protein interaction architecture. PLoS Biol. 19, 3001344; 10.1371/journal.pbio.3001344 (2021).

11. Tang, D. et al. Cryo-EM structure of C9ORF72–SMCR8–WDR41 reveals the role as a GAP for Rab8a and Rab11a. Proc. Natl. Acad. Sci. U.S.A. 117, 9876–9883; 10.1073/pnas.1916796117 (2020).

12. Khan, A. R. & Ménétrey, J. Structural biology of Arf and Rab GTPases’ effector recruitment and specificity. Structure 21, 1284–1297; 10.1016/j.str.2013.06.016 (2013).

13. DeJesus-Hernandez, M. et al. Expanded GGGGCC hexanucleotide repeat in noncoding region of C9ORF72 causes chromosome 9p-linked FTD and ALS. Neuron 72, 245–256; 10.1016/j.neuron.2011.09.011 (2011).

14. Shi, Y., et al. Haploinsufficiency leads to neurodegeneration in C9ORF72 ALS/FTD human induced motor neurons. Nat. Med. 24, 313–325; 10.1038/nm.4490 (2018).

15. Zhu, Q. et al. Reduced C9ORF72 function exacerbates gain of toxicity from ALS/FTD-causing repeat expansion in C9orf72. Nat. Neurosci. 23, 615–624; 10.1038/s41593-020-0619-5 (2020).

16. Farrell, K. B. et al. The tyrosine kinase Pyk2 regulates Arf1 activity by phosphorylation and inhibition of the Arf-GTPase-activating protein ASAP1. J. Biol. Chem. 295, 12247–12260; 10.1074/jbc.RA120.013519 (2020).

17. Su, M.-Y. et al. Structural basis for the ARF GAP activity and specificity of the C9orf72 complex. Nat. Commun. 12, 3786; 10.1038/s41467-021-24050-5 (2021).

18. Banerjee, P. et al. ProTox-II: a webserver for the prediction of toxicity of chemicals. Nucleic Acids Res. 46, W257–W263; 10.1093/nar/gky318 (2018).

19. Friesland, A. et al. Small molecule targeting Cdc42–intersectin interaction disrupts Golgi organization and suppresses cell motility. Proc. Natl. Acad. Sci. U.S.A. 110, 1261–1266; 10.1073/pnas.1215507110 (2013).

20. Aguilar, B. J. et al. Inhibition of Cdc42–intersectin interaction by small molecule ZCL367 impedes cancer cell cycle progression, proliferation, migration, and tumor growth. Cancer Biol. Ther. 20, 740–749; 10.1080/15384047.2018.1564554 (2019).

21. Ward, T. H., Polishchuk, R. S., Caplan, S., Hirschberg, K. & Lippincott-Schwartz, J. Maintenance of Golgi structure and function depends on the integrity of ER export. J. Cell Biol. 155, 557–570. 10.1083/jcb.200107045 (2001).

22. Liu, W., Duden, R., Phair, R. D. & Lippincott-Schwartz, J. ArfGAP1 dynamics and its role in COPI coat assembly on Golgi membranes of living cells. J. Cell Biol. 168, 1053–1063. 10.1083/jcb.200410142 (2005).

23. Ackema, K. B. et al. The small GTPase Arf1 modulates mitochondrial morphology and function. EMBO J. 33, 2659–2675. 10.15252/embj.201489039 (2014).

24. Enkler, L. et al. Arf1 coordinates fatty acid metabolism and mitochondrial homeostasis. Nat. Cell Biol. 25, 1157–1172. 10.1038/s41556-023-01180-2 (2023).

25. Rambold, A.S., Kostelecky B., Elia N.,& Lippincott-Schwartz J. Tubular network formation protects mitochondria from autophagosomal degradation during nutrient starvation. Proc. Natl. Acad. Sci. U.S.A. 108, 10190–10195. 10.1073/pnas.1107402108 (2011).

26. Waller, T. J., Collins, C. A., & Dus, M. Pyruvate kinase deficiency links metabolic perturbations to neurodegeneration and axonal protection. Molecular Metabolism. 98, 102187. 10.1016/j.molmet.2025.102187 (2025).

27. Haass, C. et al. Amyloid β-peptide is produced by cultured cells during normal metabolism. Nature. 359, 322–325. 10.1038/359322a0 (1992).

28. Zheng, H. & Koo, E. H. The amyloid precursor protein: beyond amyloid. Mol. Neurodegener. 1, 5. 10.1186/1750-1326-1-5 (2006).

29. Rice, H. C., Young-Pearse, T. L. & Selkoe, D. J. Systematic evaluation of candidate ligands regulating ectodomain shedding of amyloid precursor protein. Biochemistry. 52, 3264–3277. 10.1021/bi400165f (2013).

30. Lei Liu, Li Ding, Matteo Rovere, Michael S. Wolfe, Dennis J. Selkoe; A cellular complex of BACE1 and γ-secretase sequentially generates Aβ from its full-length precursor. J Cell Biol. 218, 644–663. 10.1083/jcb.201806205 (2019).

31. Matrone, C., Ciotti, M. T., Mercanti, D., Marolda, R. & Calissano, P. NGF and BDNF signaling control amyloidogenic route and Aβ production in hippocampal neurons. Proc. Natl. Acad. Sci. U.S.A. 105, 13139–13144. 10.1073/pnas.0806133105 (2008).

32. Nebenführ, A., Ritzenthaler, C. & Robinson, D. G. Brefeldin A: deciphering an enigmatic inhibitor of secretion. Plant Physiol. 130, 1102–1108; 10.1104/pp.011569 (2002).

33. Zeeh, J.-C. et al. Dual specificity of the interfacial inhibitor brefeldin A for Arf proteins and Sec7 domains. J. Biol. Chem. 281, 11805–11814; 10.1074/jbc.M600149200 (2006).

34. Viaud, J. et al. Structure-based discovery of an inhibitor of Arf activation by Sec7 domains through targeting of protein-protein complexes. Proc. Natl. Acad. Sci. U.S.A. 104, 10370–10375; 10.1073/pnas.0703592104 (2007).

35. Jansen, R. M. & Hurley, J. H. Longin domain GAP complexes in nutrient signalling, membrane traffic and neurodegeneration. FEBS Lett. 597, 750–761; 10.1002/1873-3468.14580 (2023).

36. Zhai, J. et al. Inhibition of cytohesins protects against genetic models of motor neuron disease. J. Neurosci. 35, 9088–9105; 10.1523/JNEUROSCI.0685-15.2015 (2015).

37. Loving, K., Alberts, I. & Sherman, W. Energetic analysis of fragment docking and application to structure-based pharmacophore hypothesis generation. J. Comput. Aided Mol. Des. 23, 541–554; 10.1007/s10822-009-9268-1 (2009).

38. Kikuchi, S., et al. Brefeldin A-induced neurotoxicity in cultured spinal cord neurons. J. Neurosci. Res. 71, 591–599; 10.1002/jnr.10536 (2003).

39. Franzusoff, A., Redding, K., Crosby, J., Fuller, R. S. & Schekman, R. Localization of components involved in protein transport and processing through the yeast Golgi apparatus. J. Cell Biol. 112, 27–37. 10.1083/jcb.112.1.27 (1991).

40. Richardson B, McDonold C, Fromme J. The Sec7 Arf-GEF Is Recruited to the *trans*-Golgi Network by Positive Feedback. Developmental Cell. 22, 799–810. 10.1016/j.devcel.2012.02.006 (2012).

41. B.A. Brownfield, B.C. Richardson, S.L. Halaby, J.C. Fromme, Sec7 regulatory domains scaffold autoinhibited and active conformations, Proc. Natl. Acad. Sci. U.S.A. 121. 10.1073/pnas.2318615121 (2024).

42. Belosludtseva, N.V.; Matveeva, L.A.; Belosludtsev, K.N. Mitochondrial Dyshomeostasis as an Early Hallmark and a Therapeutic Target in Amyotrophic Lateral Sclerosis. Int. J. Mol. Sci. 24, 16833. 10.3390/ijms242316833 (2023).

43. Niu, T.K.; Pfeifer, A.C.; Lippincott-Schwartz, J.; Jackson, C.L. Dynamics of GBF1, a Brefeldin A-sensitive Arf1 exchange factor at the Golgi. Mol Biol Cell. 16, 1213–1222. doi: 10.1091/mbc.e04-07-0599 (2005).

44. Madhavi Sastry, G., Adzhigirey, M., Day, T., Annabhimoju, R. & Sherman, W. Protein and ligand preparation: parameters, protocols, and influence on virtual screening enrichments. J. Comput. Aided Mol. Des. 27, 221–234; 10.1007/s10822-013-9644-8 (2013).

45. Halgren, T. A. Identifying and characterizing binding sites and assessing druggability. J. Chem. Inf. Model. 49, 377–389; 10.1021/ci800324m (2009).

46. Kenyon, V. et al. Novel human lipoxygenase inhibitors discovered using virtual screening with homology models. J. Med. Chem. 49, 1356–1363; 10.1021/jm050934s (2006).

47. Friesner, R. A. et al. Extra precision Glide: docking and scoring incorporating a model of hydrophobic enclosure for protein-ligand complexes. J. Med. Chem. 49, 6177–6196; 10.1021/jm051256o (2006).

48. Pattar, S. V. et al. In silico molecular docking studies and MM/GBSA analysis of coumarin-carbonodithioate hybrid derivatives divulge the anticancer potential against breast cancer. Beni-Suef Univ. J. Basic Appl. Sci. 9, 36; 10.1186/s43088-020-00060-6 (2020).

49. Amick, J. et al. C9orf72 binds SMCR8, localizes to lysosomes, and regulates mTORC1 signaling. Mol. Biol. Cell 27, 3040–3051; 10.1091/mbc.E16-01-0004 (2016).

