## Supplemental Material for "Small molecules targeting ARF1 interaction with C9orf72:SMCR8:WDR41 complexes suppress its overactivation implicated in ALS/FTD"

*: Corresponding author:

**Supplementary Table S1. Docking scores, binding free energy (MM/GBSA), and interaction patterns of top ranking SMMs targeting ARF1-CSW interactions**

| Compounds | Vina dock Score (kcal/mol) | Dock Score (kcal/mol) | Binding Energy (kcal/mol) | Interactions |
|---|---|---|---|---|
| MCULE-5095997944 | -8.1 | -5.37 | -60.89 | Hydrogen bonds: ILE49(E), PRO107(B).<br>Halogen bond: ARG151(B).<br>Hydrophobic interactions: PHE51(E), PRO152 (B), VAL150(B), HIS138(B).<br>Π-π interactions: HIE65(C). |
| MCULE-2964877735 | -7.9 | -6.45 | -59.27 | Hydrogen bonds: HIE106(B), SER61(B), ARG151(B).<br>Hydrophobic interactions: PHE149(B), PRO152 (B), VAL150(B), CYS154(B), ILE46(E), ILE49(E). |
| MCULE-3711620516 | -7.8 | -6.39 | -56.63 | Hydrogen bonds: VAL53(E), HIE65(C), GLY50 (E), ILE49(E).<br>Hydrophobic interactions: SER110(B), GLY109(B), PRO107(B), ASN6G4(C), ASN68(C), TYR32(C). |
| MCULE-2336465708 | -7.6 | -4.67 | -54.29 | Hydrogen bonds: PRO107(B), ILE49(E).<br>Hydrophobic interactions: PHE149(B), VAL150(B), PRO152(B), VAL136(B), PRO108(B), ILE46(E), PHE51(E). |
| MCULE-9770532735 | -7.6 | -4.92 | -41.38 | Hydrogen bonds: GLY109(B).<br>Hydrophobic interactions: PHE149(B), VAL150(B), PRO152(B), ARG151(B), ILE49(E), ILE46(E), PRO107(B). |
| MCULE-8081431558 | -7.4 | -5.73 | -61.76 | Hydrogen bonds: GLY109(B).<br>Hydrophobic interactions: PHE149(B), VAL150(B), PRO152(B), ARG151(B), ILE49(E), ILE46(E), PRO107(B). |

| | | | | |
|---|---|---|---|---|
| MCULE-1582781505 | -7.4 | -6.24 | -61.06 | Hydrogen bonds: GLY109(B), GLY50(E), PHE51(E).<br><br>Hydrophobic interactions: ILE46(E), PRO47(E), ILE49(E), VAL150(B),<br><br>Π-π interactions: PHE149(B). |
| MCULE-5055852153 | -7.1 | -4.55 | -60.61 | Hydrogen bonds: GLY109(B), ARG151(B).<br><br>Hydrophobic interactions: PHE149(B), VAL150(B), PRO152(B), PRO(108), PRO107(B), TYR35(C), ILE49(E), ILE46(E), VAL43(E). |
| MCULE-9316017353 | -6.9 | -5.55 | -59.58 | Hydrogen bonds: VAL53(E), PHE51(E), GLY50(E), HIE65(E).<br><br>Hydrophobic interactions: PRO107(B), ILE46(E), PHE51(E), VAL53(E). |
| MCULE-7957768027 | -6.8 | -6.22 | -71.41 | Hydrogen bonds: GLY50(E), PHE51(E), ILE49(E), GLY109(B), ARG151(B).<br><br>Hydrophobic interactions: PHE149(B), VAL150(B), PRO108(B), PRO152(B)<br><br>Π-π interactions: HIE65(C) |
| MCULE-7224174737 | -6.5 | -4.65 | -52.3 | Hydrogen bonds: GLY109(B), ARG151(B), ASN52(E).<br><br>Hydrophobic interactions: PHE149(B), VAL150(B), PRO108(B), PRO152(B), PHE61(C), ILE49(E), ILE46(E). |
| MCULE-6491018430 | -6.4 | -4.08 | -60.86 | Hydrogen bonds: GLY109(B), ILE49(E), HIE65(C).<br><br>Hydrophobic interactions: PRO107(B), PRO108(B), PHE51(E), VAL53(E), PHE61(C), VAL150(B), VAL136(B), TYR32(C). |
| MCULE-5145044294 | -6.3 | -5.63 | -57.4 | Hydrogen bonds: HIE65(C), ILE49(E).<br><br>Hydrophobic interactions: PHE51(E), ILE46(E), PRO108(B), PRO107(B), PHE61(C).<br><br>Π-π interactions: HIE65(C) |

Note: Compounds were docked into binding pockets of ARF1-CSW complex using the XP mode of Glide module in Schrodinger with the default settings. Docking scores and detailed interactions with different proteins in interface of ARF1 and CSW complex of are listed. Chains E, B and C are related to ARF1, C9orf72 and SMCR8 respectively.

### Supplementary Table S2. Physicochemical properties of the representative initial hits

| Properties | parameter | MCULE-5095997944 | MCULE-2964877735 | MCULE-3711620516 | MCULE-5145044294 | MCULE-5055852153 | MCULE-2336465708 |
|---|---|---|---|---|---|---|---|
| Physico-chemical properties[a] | MW(g/mol) | 368.77 | 321.283 | 352.30 | 337.38 | 323.354 | 335.365 |
| | Number of atoms | 26 | 23 | 26 | 25 | 24 | 25 |
| | Rotatable bonds | 5 | 2 | 4 | 5 | 6 | 3 |
| | H-bond acceptor | 6 | 6.5 | 9.5 | 6.5 | 6.5 | 7.7 |
| | H-bond donors | 1.5 | 2 | 1 | 1 | 1 | 1 |
| | Molar Refractivity | 99.87 | 80.53 | 93.59 | 94.61 | 89.81 | 97.30 |
| | Volume | 1123.12 | 944.27 | 1069.13 | 1116.56 | 1053.11 | 1060.49 |
| Aqueous solubility [b] | QPlogS | -5.81 | -3.78 | -4.03 | -4.85 | -4.23 | -4.41 |
| Lipinski RO5 | No. of Violations | 0 | 0 | 0 | 0 | 0 | 0 |

[a]The physico-chemical properties of initial hits (MCULE-5095997944, MCULE-2964877735, MCULE-5145044294, MCULE-5055852153 and MCULE-2336465708).

[b] Aqueous solubility of a molecule in log mol/L units (the recommended range of QplogS for drug-like molecules is -6.5 to 0.5).

[c] Number of violations of Lipinski Rule of Five.

**Supplementary Table S3.** ***In silico*** **ADME assessment of the representative initial hits**

| Compound ID | | MCULE-5095997944 | MCULE-2964877735 | MCULE-3711620516 | MCULE-5145044294 | MCULE-5055852153 | MCULE-2336465708 |
|---|---|---|---|---|---|---|---|
| **Absorption** | **PSA[a]** | 128.286 | 107.09 | 150.92 | 83.13 | 83.101 | 104.59 |
| | **logPo/w[b]** | 3.05 | 1.54 | 0.54 | 2.74 | 2.088 | 3.23 |
| | **logPw[c]** | 12.36 | 12.10 | 14.90 | 10.92 | 12.747 | 14.37 |
| | **Percentage oral absorption[d]** | 81.98 | 77.98 | 57.52 | 96.24 | 93.22 | 90.59 |
| **Distribution** | **QPlogBB[e]** | -1.79 | -1.09 | -2.37 | -0.73 | -0.592 | -1.23 |
| | **SASA[f]** | 672.67 | 554.29 | 631.13 | 606.95 | 617.65 | 656.93 |
| | **FOSA[g]** | 12.98 | 196.09 | 83.3 | 201.97 | 186.41 | 23.59 |
| | **QplogKhsa[h]** | 0.2 | -4.03 | -4.91 | 0.046 | -0.092 | 0.18 |
| **Metabolism and Excretion** | **QPPMDCK[i]** | 122.47 | 164.86 | 12.72 | 467.04 | 519.72 | 351.06 |
| | **QPPcaco[j]** | 118.95 | 221.35 | 33.83 | 948.14 | 1046.70 | 315.33 |

The QikProp tool of Schrödinger was used for prediction of pharmacokinetic properties criteria of initial hits (MCULE-5095997944, MCULE-2964877735, MCULE-5145044294, MCULE-5055852153 and MCULE-2336465708).

[a] Van der Waals surface area of polar nitrogen and oxygen atoms and carbonyl carbon atoms. (7.0 to 200.0).

[b] Predicted water/gas partition coefficient. (4.0 to 45.0).

[c] Predicted octanol/water partition coefficient. (–2.0 to 6.5).

[d] the brain/blood partition coefficient (-3.0 to 1.2).

[e] Predicted human oral absorption (on 0 to 100% scale. >80% is high, <25% is poor).

[f] Total solvent accessible surface area (SASA) in square angstroms using a probe with a 1.4 Å radius. (300.0 to 1000.0).

[g] Hydrophobic components of the SASA (saturated carbon and attached hydrogen). (0.0 to 750.0).

[h] Prediction of binding to human serum albumin. (–1.5 to 1.5).

[i] Predicted apparent MDCK cell permeability in nm/sec. (<25 poor, >500 great QikProp predictions are for non-active transport. <25 poor, >500 great).

[j] Predicted apparent Caco-2 cell permeability in nm/sec. Caco2 cells are a model for the gut-blood barrier.

### Supplementary Table S4. Toxicity properties of top hit compound SCC944

| Endpoint | Target | MCULE-5095997944 (SCC944) |
|---|---|---|
| **Organ toxicity** | **Hepatotoxicity** | Active (0.59) |
| | **Neurotoxicity** | Active (0.68) |
| | **Nephrotoxicity** | Inactive (0.55) |
| | **Respiratory toxicity** | Active (0.52) |
| | **Cardiotoxicity** | Inactive (0.82) |
| **Toxicity endpoints** | **Carcinogenicity** | Inactive (0.52) |
| | **Immunotoxicity** | Inactive (0.98) |
| | **Mutagenicity** | Inactive (0.56) |
| | **Cytotoxicity** | Inactive (0.73) |
| | **Ecotoxicity** | Inactive (0.71) |
| | **Toxicity Class** | 5 |
| | **Nutritional toxicity** | Inactive (0.67) |
| **Tox21-nuclear receptor signaling pathway** | **Androgen Receptor (AR)** | Inactive (0.88) |
| | **Aryl hydrocarbon receptor (AhR)** | Inactive (0.5) |
| **Tox21-stress response pathway** | **Heat shock factor response element** | Inactive (0.91) |
| **Predict lethal dose** | **LD50 (mg/Kg)** | 3000 |

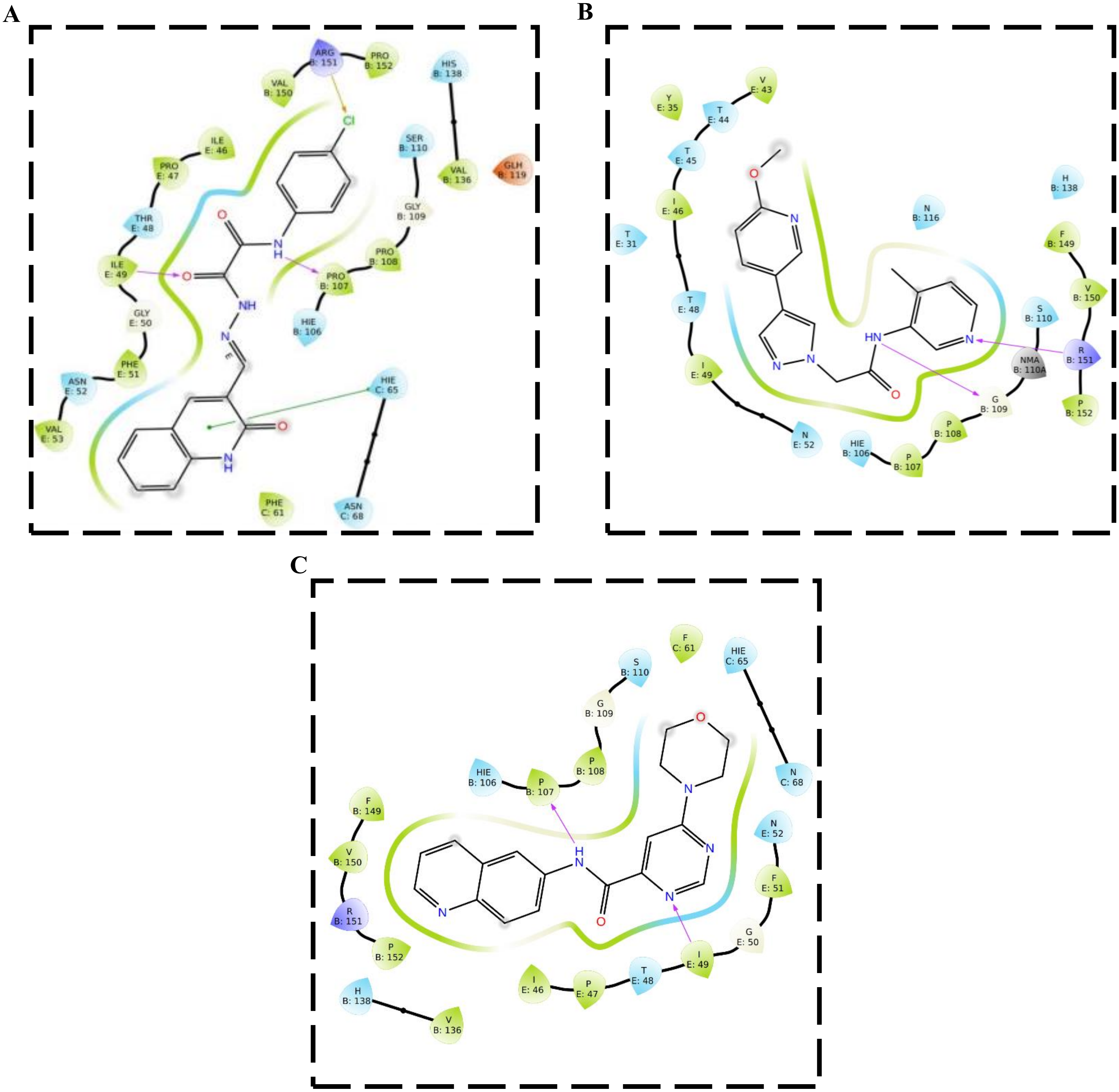


**Supplementary Figure S1. 2D illustration of initial hit compound interactions into ARF1-CSW complex.** 2D representation of binding site residues of ARF1-CSW complex proteins with (**A**) **SCC944**, (**B**) **SCC153** and (**C**) **SCC708**. The hydrogen bonds, hydrophobic, polar interactions, charged negative, charged positive are represented by purple, green, blue, orange, and violet color arrows, respectively.
